# Reversible fixation of microbial cells for targeted genomics

**DOI:** 10.64898/2026.10.12.758561

**Authors:** J. Pamela Engelberts, Kenneth Wasmund, Marc Mussman, Nataliia Solntseva, Fatima C. Pereira, Christian Rinke, Apoorva Prabhu, James G. Volmer, Buck T. Hanson, Bela Hausmann, Jay Osvatic, Gene W. Tyson, Simon J. McIlroy, Alexander Loy

## Abstract

Phylogeny- or activity-based fluorescent labelling of microorganisms coupled with fluorescence activated cell sorting (FACS) is a powerful approach to rapidly isolate cells of interest from complex communities for targeted genomics. However, the method has not been widely adopted because standard paraformaldehyde or ethanol fixation procedures hamper nucleic acid recovery and/or long-term sample storage. Here, we present dithiobis(succinimidyl propionate) (DSP) as a reversible fixative for microbial cells that enables fluorescence imaging and DNA extraction and amplification for sequencing. DSP fixation is compatible with widely used fluorescence *in situ* hybridization (FISH) approaches and bioorthogonal non-canonical amino acid tagging, and can be applied to diverse sample types, where it consistently performed as well as paraformaldehyde. Furthermore, high-quality genomes were obtained from individual and multiple bacterial cells selectively extracted from complex samples using FISH-FACS. Thus, DSP is a dual-use fixative for microbiome imaging and sequencing, improving targeted microbial genomics.

## Introduction

Metagenomic sequencing is a powerful approach to recover genomic material of microorganisms without cultivation, providing key insights into microbial community composition, metabolic potential, and evolution ^1^. However, standard metagenomic approaches still struggle to capture strain-level heterogeneity, recover rare microbial species and strains, and link mobile genetic elements (e.g., phages and/or plasmids) to their hosts ^2^. One method to overcome these limitations is single cell genomics (SCG) ^3^. However, SCG is often based on the random collection of cells, hampering the recovery of genomes of microorganisms of interest. Therefore, targeted approaches that physically extract microbes of interest from complex microbial communities prior to sequencing are increasingly being explored ^4^.

Individual microorganisms can be selectively sorted from complex communities by various parameters, including cell size and ‘granularity’, Raman-spectra of specific cellular compounds and/or isotopic labels, or fluorescence. Fluorescence can be intrinsic, derived from phylogenetically targeted probe-based labelling approaches (e.g., fluorescence *in situ* hybridization, FISH ^5^; catalysed reporter deposition-FISH, CARD-FISH ^6^; GenomeFISH ^7^), or linked to cellular activity through metabolic labelling techniques (e.g., bioorthogonal non-canonical amino acid tagging, BONCAT ^8^ and substrate analogue probing ^9^). The application of FISH to physically extract microbes is especially powerful because rRNA-targeted oligonucleotide probes or whole genome probes can be used to target microorganisms or groups of interest, from higher taxonomic levels down to strain-level. Individual FISH-stained cells can subsequently be sorted in high-throughput via fluorescence activated cell sorting (FACS), making FISH-FACS a highly efficient means to collect microbes of interest for subsequent DNA amplification, genome sequencing, and analyses ^10^. However, these methods have seen limited use, as standard cell fixation procedures for FISH hamper the recovery and enzymatic amplification of genomic DNA after cell sorting/extraction ^11^.

Microbial cells are typically fixed with paraformaldehyde (PFA) or ethanol to preserve rRNA and cellular morphology to enable access of FISH probes into cells and obtain detectable fluorescent signal intensities ^12,13^. However, ethanol significantly reduces the coverage of recovered single amplified genomes (SAGs) and PFA prevents genome amplification by cross- linking DNA ^11,14^. While formaldehyde cross-links can be reversed by chemical treatments or extended heating, it damages and chemically modifies DNA ^15^. One method that avoids fixation and has been successfully combined with FACS is fixation-free FISH ^14^. However, fixation-free FISH generally results in poor detection rates with environmental samples and requires that cells are collected fresh for hybridisation. To enable targeted genomics, a fixative that preserves microbial cells without compromising gene or genome amplification and sequencing is needed ^2^.

Here, we introduce dithiobis(succinimidyl propionate) (DSP) as a functional fixative for bacterial and archaeal cells that enables downstream DNA sequencing. DSP is a protein-specific cross-linker that is used for the preservation of eukaryotic cells for RNA sequencing and protein profiling ^16–18^. DSP cross-links to primary amines of amino acids and includes a disulfide bond, which can be cleaved by reducing agents such as dithiothreitol (DTT), meaning that the cross- linking can be “reversed” ^18^. Conveniently, DTT is present in the lysis buffers used for the multiple displacement amplification (MDA) of single cell genomes. We therefore reasoned that DSP may also act as an effective fixative for microbial cells by cross-linking proteins, ribosomes and other cellular structures, while also enabling DNA amplification and sequencing after cell sorting and lysis.

In this study, we tested the application of DSP for fluorescence imaging and targeted microbial genomics. We show that DSP fixation preserves microbial cell mass over several months of storage, is compatible with FISH, and results in fluorescent signals comparable to PFA fixation for isolates and environmental samples. DSP can be combined with widely used phylogenetic (e.g., FISH, CARD-FISH, GenomeFISH) and cellular activity-based (e.g., BONCAT) fluorescence labelling approaches. When combined with FACS, we show that high-quality metagenome-assembled genomes (MAGs) and SAGs of target taxa can be recovered from DSP fixed samples. Therefore, we envisage DSP will become a widely used fixative for fluorescence-based imaging, enrichment, and genome sequencing of microorganisms of interest from complex microbial communities.

## Results and discussion

### DSP is an effective fixative for storage and imaging of pure cultures and environmental samples

To evaluate DSP as a fixative for microbial cells, we compared it against the commonly used fixatives PFA and ethanol across several applications, including pure cultures and environmental samples. Slight variations in the DSP fixation method were tested among experiments and laboratories, differing mainly in the number of DSP additions to the samples (i.e., x1, x2 or x3 as indicated in parentheses throughout). A recommended DSP-fixation protocol can be found in the Supplementary Information section.

To evaluate the effectiveness of DSP as a fixative for microbial cells for standard FISH, we fixed stationary-phase cells from pure cultures of *Escherichia coli*, *Bacteroides thetaiotaomicron* and *Bacillus subtilis* with PFA, DSP (x1), DSP (x3), or ethanol, and stored them for 1.5, 2, and 3.5 months. We then performed FISH with a FITC mono-labelled EUB338-I ^19^ probe and evaluated the average fluorescence intensity of hybridised cells (**Table S1**). The fluorescence intensity of FISH hybridised cells of *E. coli* and *B. thetaiotaomicron* was similar among DSP and PFA treatments over the three time points, with ethanol-fixed cells generally having significantly lower fluorescence intensity than PFA- and DSP-fixed cells (Kruskal-Wallace p = <0.05) (**Fig. 1a,b and S1a**). Average fluorescence intensity of DSP (x1)-fixed *E. coli* cells was significantly higher than that of ethanol-fixed cells after 2 months of cell storage (Kruskal-Wallace p = <0.05) (**Fig. 1a**), and DSP (x3)-fixed *E. coli* cells showed significantly higher average fluorescence intensity than ethanol-fixed cells after 2 and 3.5 months (Kruskal-Wallace p = <0.05) (**Fig. 1a**). After 1.5 months of storage, the average fluorescence intensity of DSP-fixed *E. coli* cells was also noticeably higher, although not statistically significant (**Fig. 1a**). Average fluorescence intensity of DSP-fixed *B. thetaiotaomicron* cells was significantly higher than ethanol-fixed cells after all three periods of cell storage (Kruskal-Wallace p = <0.05) (**Fig. 1b and S1b**). Average fluorescence intensities of DSP (x1)-fixed and DSP (x3)-fixed cells were not significantly different across most comparisons. In contrast to *E. coli* and *B. thetaiotaomicron*, cells of *B. subtilis* gave poor or no fluorescent signals when fixed with DSP or PFA (results not shown). The poor performance of PFA fixation was expected, as PFA cross-linking of the cell wall is known to hinder probe penetration into Gram-positive bacteria ^20^. For this reason, ethanol fixation combined with sample- and taxon-specific enzymatic pretreatment is commonly recommended for FISH targeting bacteria with Gram-positive cell walls. Although DSP fixation appears to be as poor as PFA fixation for FISH of the Gram-positive *B. subtilis* cells, subsequent FISH experiments on DSP-fixed fecal samples successfully targeted other Gram-positive populations (see below). This suggests that at least some Gram-positive taxa are amenable to FISH after DSP fixation. Together, these results indicate that DSP is as effective as PFA for fixation of Gram-negative bacterial cells, is significantly more effective than ethanol, and preserves cells in a FISH compatible state for at least several months of storage.

**Figure 1.**
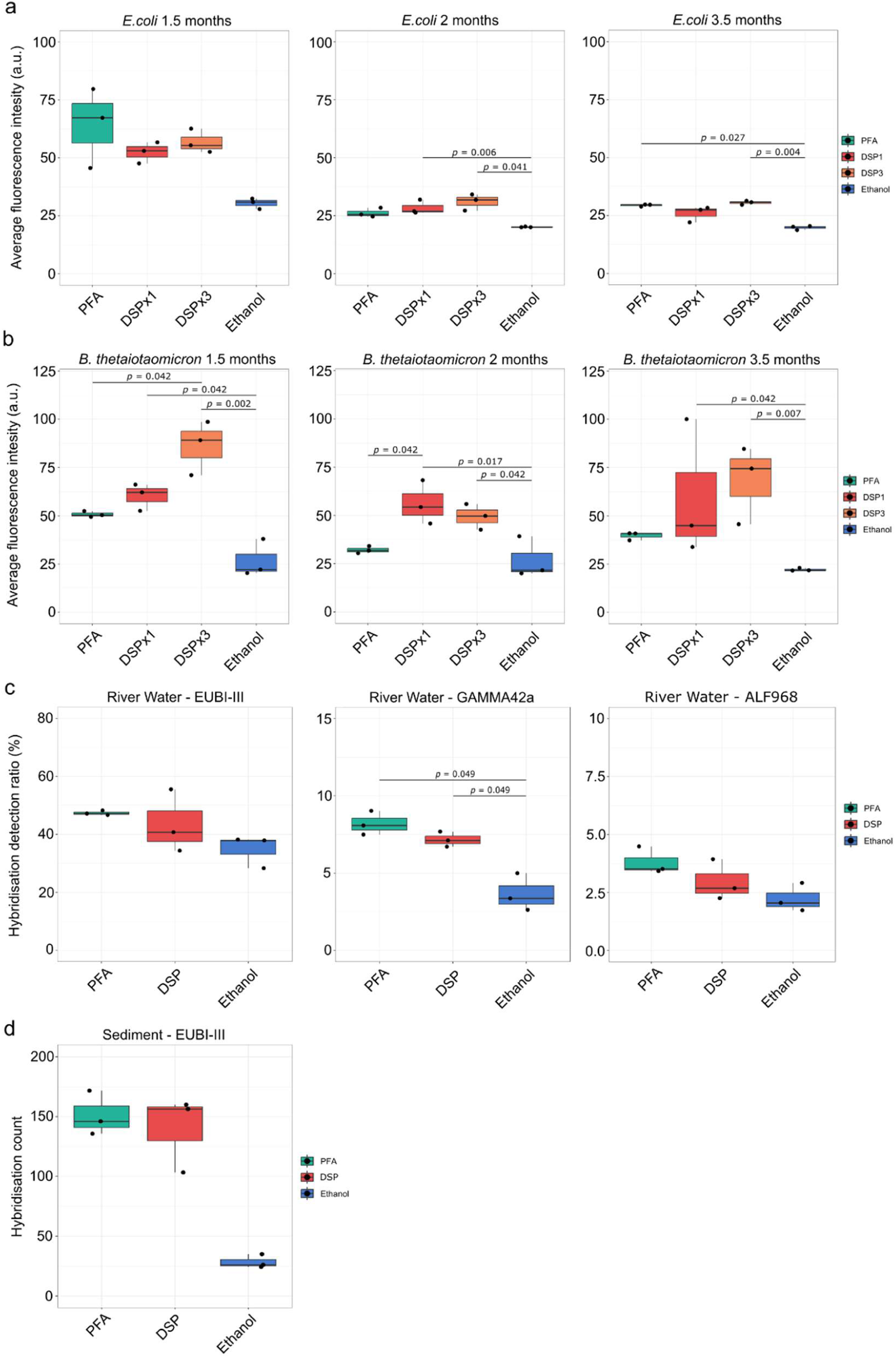
DSP is an effective fixative of bacterial cells for FISH and CARD-FISH approaches. **a**, Average Fluorescence Intensity (a.u., arbitrary units) of *E. coli* cells from FISH using FITC mono-labelled EUB338-I probe and fixed with either PFA, DSP (x1), DSP (x3), or ethanol. Cells were hybridised and imaged after 1.5, 2, and 3.5 months of storage. **b**, Average Fluorescence Intensity (A.U., arbitrary units) of *B. thetaiotaomicron* cells from FISH using FITC mono-labelled EUB338-I probe and fixed with either PFA, DSP (x1), DSP (x3), or ethanol. Cells were hybridised and imaged after 1.5, 2, and 3.5 months of storage. **c**, Hybridisation detection rates of cells from river water by CARD-FISH using probes EUB338-I-II-III probe mix targeting most Bacteria, probe GAM42a for Gammaproteobacteria, and probe ALF968 for Alphaproteobacteria. Hybridisation detection rates were determined in relation to total DAPI counts. **d**, Hybridisation detection counts of cells from marine sediments by CARD-FISH using probes EUB338-I-II-III probe mix targeting most Bacteria, showing average count per field of view for each triplicate hybridisation. Hybridisation detection counts were determined without relation to total DAPI counts because DAPI counts and signals were too low for reliable quantification from ethanol-fixed samples. For analysis of FISH experiments with pure cultures of *E. coli* and *B. thetaiotaomicron*, statistics were performed between treatments for each time point and not across all time points, to avoid differences caused by inter- experiment variation from FISH. For *E. coli* comparisons (panel a), an average of 1,423 ± 358 cells (DAPI stained) were evaluated for each treatment. For *B. thetaiotaomicron* comparisons (panel b), an average of 4,692 ± 1,160 cells (DAPI stained) were evaluated for each treatment. For river water CARD-FISH comparison with EUB-I-II-III probes (panel c), an average of 4,692 ± 1,460 cells (DAPI stained) were evaluated for each treatment, for probe GAMMA42a 7,057 ± 608 cells (DAPI stained) were evaluated for each treatment, and for probe ALF968 6,218 ± 730 cells (DAPI stained) were evaluated for each treatment. For sediment CARD-FISH comparison with EUB-I-II-III probes (panel d), an average of 957 ± 500 cells (CARD-FISH stained) were counted for each treatment. Box-and-whisker plots show the median and interquartile range; whiskers extend to values within 1.5x the interquartile range. In all box-and-whisker plots, each black-filled circle shows data points for the average values from three fields of view, for each of the triplicate hybridisations. Significance values between treatments are displayed above horizontal bars.

To assess the effectiveness of DSP as a fixative for microbial cells in environmental samples, we fixed samples from river water (Danube River, Vienna, Austria), and from estuarine marine sediment (Aveiro, Portugal) with PFA, DSP (x1 and x3), or ethanol. As environmental samples like river water and sediments are particularly challenging sample types for FISH due to autofluorescent organic and inorganic particles, and low cellular activity/ribosome content of resident microorganisms ^21^, we applied CARD-FISH to increase signal intensity ^22^. For river water samples, we performed CARD-FISH 10 days after fixation and cell storage using the EUB338-I- II-III probe mix targeting most bacteria ^19,23^, probe GAM42a for Gammaproteobacteria ^24^, and probe ALF968 for Alphaproteobacteria ^25^ (**Table S1**). Because CARD-FISH amplifies fluorescent signals ^6^ and the signals are therefore not proportionate with ribosomal RNA content, results were evaluated based on hybridization detection ratios compared to DAPI cell counts. Mean detection ratios of cells hybridised with probe EUB338-I-II-III, GAM42a, or ALF968 were slightly higher for PFA- than DSP-fixed cells, but not significantly different (**Fig. 1c and S1c**). Detection ratios of ethanol-fixed cells were generally lower than PFA- and DSP-fixed cells for probes EUB338-I-II-III and ALF968, and significantly lower for probe GAM42a hybridised cells (Kruskal-Wallace p = <0.05) (**Fig. 1c**).

For sediments, we performed CARD-FISH using the probe EUB338-I-II-III on cells that were separated from the sediment and stored for 3 days. Detected numbers of EUB338-I-II-III probe hybridised cells were at least five-fold higher for PFA- and DSP-fixed cells compared to ethanol-fixed cells, although not statistically significant (**Fig. 1d and S1d**). Detected numbers of CARD-FISH positive cells were comparable between DSP- and PFA-fixed samples (**Fig. 1d**), showing that DSP is an effective fixative for complex environmental samples like sediments.

To evaluate the compatibility of DSP with other hybridization-based labelling approaches, we fixed a human fecal sample with DSP and applied GenomeFISH. In contrast to rRNA-targeted FISH and CARD-FISH probes, GenomeFISH probes are generated by fragmenting and chemical fluorescence labelling of genomic DNA of a microorganism of interest, enabling strain-level differentiation ^7^. Genomic DNA is derived from either a pure culture or an amplified genome of a single microbial cell. Here, we generated GenomeFISH probes from the genome of an in-house cultured strain of *Agathobacter rectalis* (Gram-positive; sourced from the Australian Human Microbiome Biobank) and applied them to a human fecal sample fixed with DSP (x3). This showed that DSP fixation is compatible with GenomeFISH, obtaining signals consistent with what was previously observed in PFA fixed samples (**Fig. S2)** ^7^. It further indicated that some Gram-positive taxa are amenable to FISH after DSP fixation, which may reflect a more permeable cell wall architecture than is typical of Gram-positive cells ^26^.

To further test how DSP affects microbial cell integrity beyond fluorescence imaging methods, we performed single-cell buoyant mass measurements on human fecal microbiota. In a first experiment, fecal samples from two individuals were either left unfixed or fixed with PFA, DSP (x2), or ethanol, then stored for 1 or 42 days prior to analysis using a suspended microchannel resonator system (LifeScale, Affinity Biosensors) ^27^. For both individuals, DSP- and PFA-fixed samples exhibited single-cell buoyant mass distributions comparable to the unfixed condition at Day 1 (**Fig. 2a**). In contrast, ethanol-fixed samples showed significantly lower median masses already at Day 1, indicating substantial mass loss that persisted to Day 42. Importantly, PFA- and DSP-fixed samples did not exhibit measurable mass loss during storage up to 42 days. In a second experiment that directly compared single-cell buoyant mass of cells from PFA- and DSP-fixed samples from across an additional 30 fecal specimens, pairwise comparisons demonstrated a strong linear correlation and no statistically significant differences in median single-cell buoyant mass (**Fig. 2b,c**). Consistent with these findings, previous analyses comparing unfixed and PFA-fixed cells from pure cultures of *Escherichia coli* and *Vibrio cyclitrophicus* reported only minor effects of fixation on single-cell dry mass ^27^. Overall, both PFA and DSP fixation preserved the buoyant mass and thus presumably the cellular content of individual microbial cells in a natural sample effectively, even after prolonged storage for six weeks. In contrast, ethanol fixation led to substantial and persistent mass loss, which is consistent with the proposed decrease of structural integrity of some bacterial cell types with alcohol-based fixatives^28^.

**Figure 2.**
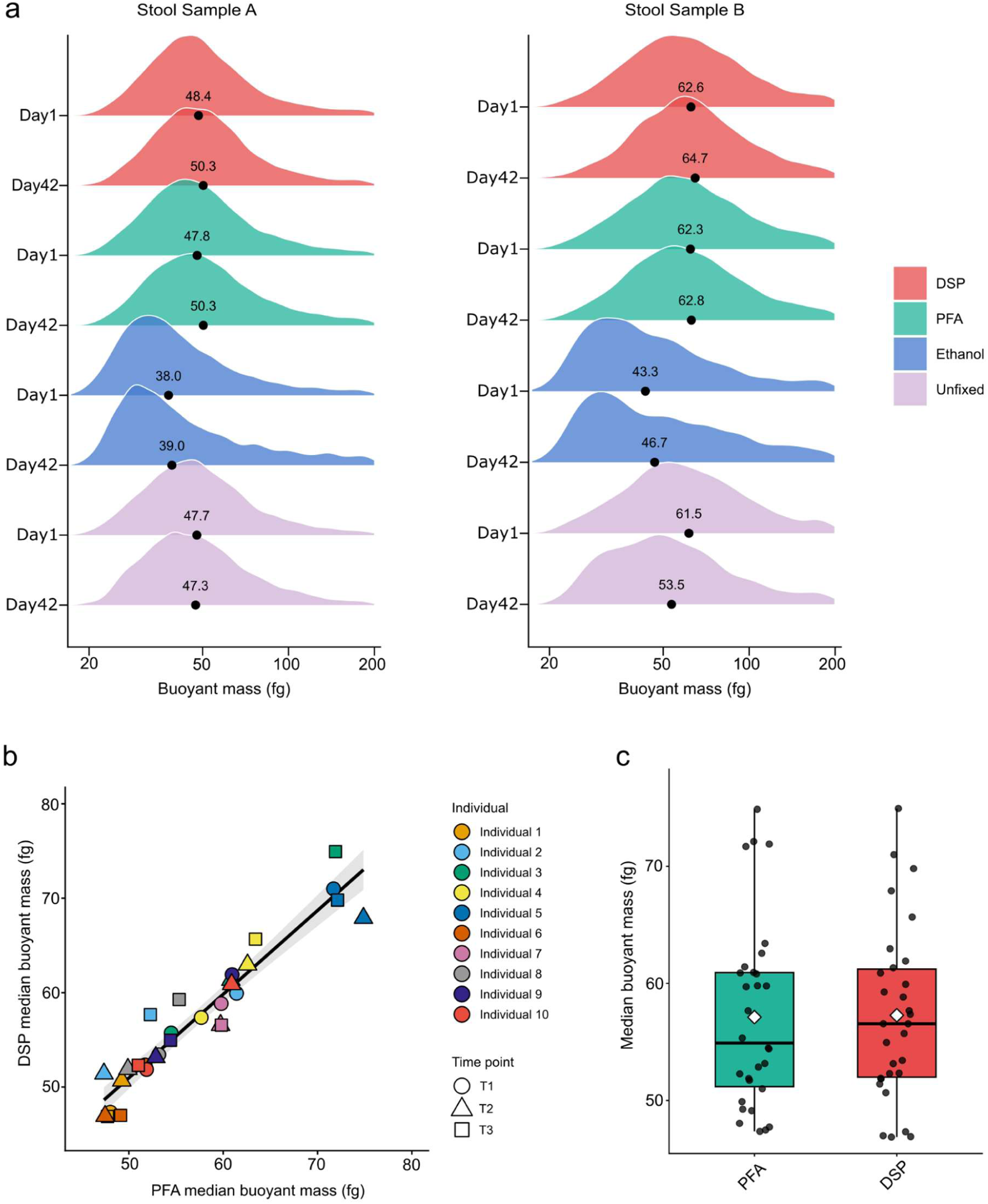
PFA and DSP fixation equally preserve single-cell buoyant mass in human fecal microbiota. Distribution patterns and median values are based on buoyant mass measurements of 2,236 to 46,500 cells per sample (average DSP = 7,346: PFA = 10,652). **a,** Single-cell buoyant mass distributions in two stool samples A and B that were either left unfixed or fixed with PFA, DSP (x2), or ethanol and then stored for 1 and 42 days. At day 1, DSP- and PFA-fixed stool samples showed single-cell buoyant mass distributions comparable to unfixed controls (medians ≈48 fg for Sample A and ≈62 fg for Sample B). In contrast, ethanol fixation produced significantly lighter cells at day 1 (≈38 fg and ≈43 fg, respectively), with reduced masses persisting to day 42. Over ∼6 weeks of storage, DSP- and PFA-fixed samples remained largely stable, while unfixed Sample B showed a modest mass decrease (from 62 to 54 fg). **b**, Pairwise comparison of median single-cell buoyant mass in PFA- and DSP-fixed feces (n = 30 fecal samples; ten individuals, three time points each) showed high linear correlation (Spearman’s ρ = 0.962, p < 2.2x10⁻¹⁶, grey shading indicates the 95% confidence for the fitted regression mean lm(DSP ∼ PFA) shown as black line). **c**, Box-and-whisker plot of the data from panel **b**, showing no difference between DSP and PFA (paired Wilcoxon signed-rank test, p = 0.666). White diamonds show the mean of sample medians (DSP: 57.3 fg; PFA: 57.1 fg). Box-and-whisker plots show the median and interquartile range; whiskers extend to values within 1.5x the interquartile range. Each black-filled circle shows the median single-cell buoyant mass of all cells measured in each sample.

Together, these results highlight that DSP is an effective fixative for imaging of pure cultures, environmental samples and human samples, and can be combined with widely applied labelling approaches (e.g., FISH, CARD-FISH, and GenomeFISH). It further showed that DSP fixation can maintain cells amenable for FISH over at least several months of storage and maintains the integrity of microbial cells similar to PFA fixation.

### DSP fixation enables the recovery of 16S rRNA gene amplicon sequences, MAGs, and SAGs of targeted taxa

A major benefit of DSP is the ability to reverse the fixation with reducing agents (e.g., DTT), enabling the downstream amplification and sequencing of DNA ^18^. To verify that DNA can be recovered from targeted microbial cells, we first investigated the compatibility of DSP fixation with 16S rRNA gene amplicon sequencing (using BONCAT-FACS), after which we explored the ability to recover MAGs and SAGs from targeted populations through mini-metagenomes and single-cell genomics using FISH-FACS.

### DSP fixation is effective for BONCAT-FACS and 16S rRNA gene amplicon sequencing

BONCAT is commonly used to assess the translational activity of microbes at the single-cell level by fluorescence-tagging of newly synthesized proteins ^8,29–32^. BONCAT can be coupled with FACS for downstream sequencing of sorted cells, which is typically performed with unfixed or ethanol-fixed samples ^29,32–34^. Here, we tested DSP fixation for BONCAT-FACS of fecal samples. We incubated a human fecal sample from a single donor in a nutrient-limited medium supplemented with either glucose, trehalose or no substrate addition as control in the presence of L-azidohomoalanine (AHA), then fixed samples after 4 and 6 hrs of incubation with PFA, DSP (x3), or ethanol. After 3 and 12 days of cell storage, we performed click reactions with fluorescently-labelled alkyne probes, then evaluated relative detection rates of BONCAT positive cells versus total DAPI counts. This showed that DSP fixation yielded BONCAT-positive detection rates significantly higher than ethanol-fixed samples in 8 of 12 comparisons (**Fig. 3a,b and S3**). It also showed that DSP fixation yielded BONCAT-positive detection rates that were statistically not different to PFA for all 12 comparisons (**Fig. 3a,b and S3**).

**Figure 3.**
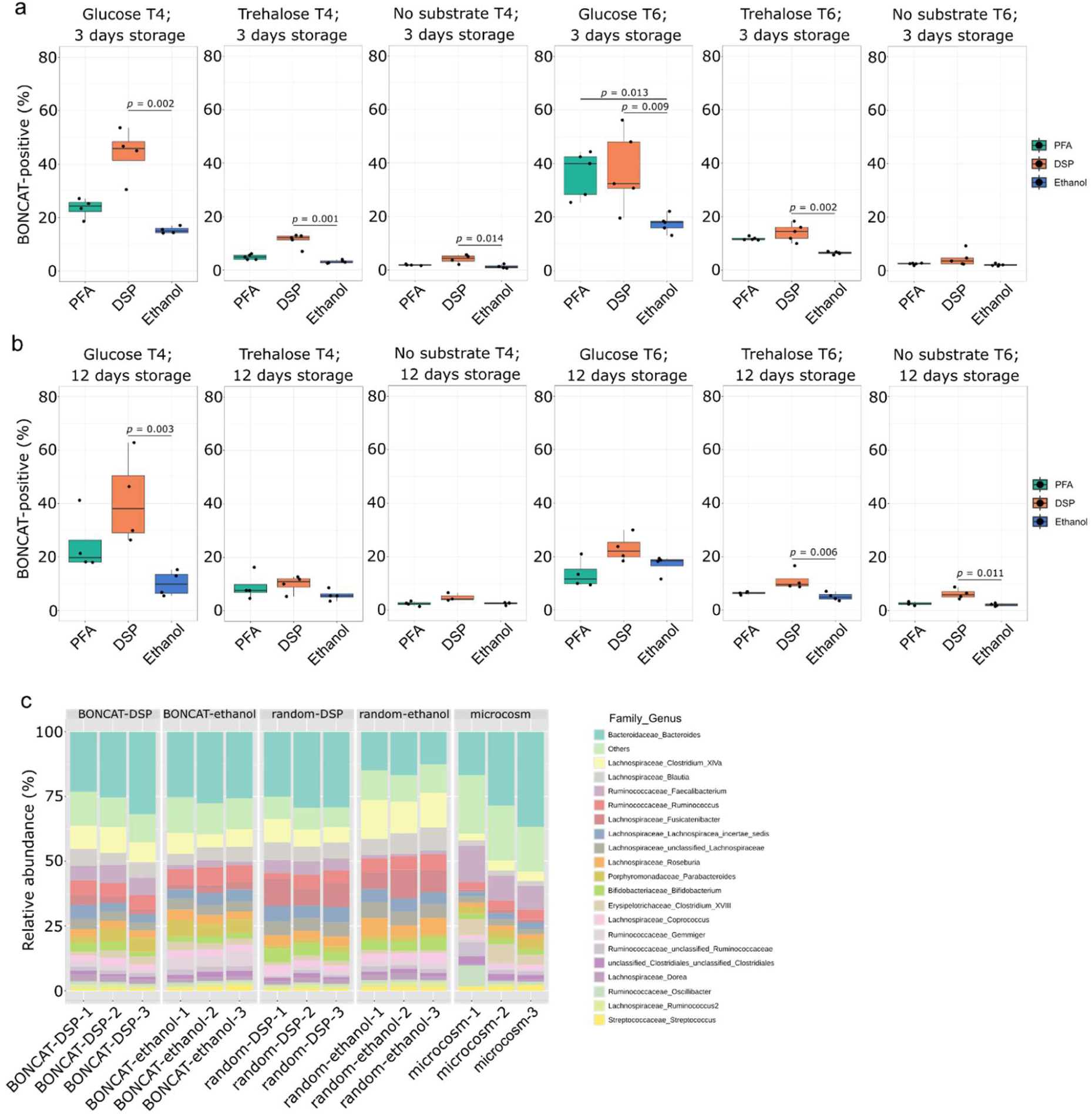
DSP fixation is compatible with BONCAT imaging and downstream PCR and amplicon sequencing. **a**, BONCAT-positive (Cy5) detection ratios (%) relative to DAPI-stained cell counts for glucose and trehalose amended fecal microcosms, which were fixed with either PFA, DSP (x3) or ethanol after 4 (T4) or 6 hrs (T6) of incubation, and stored for 3 days before click reactions and imaging. **b**, BONCAT-positive (Cy5) detection ratios (%) relative to DAPI-stained cell counts for glucose and trehalose amended fecal microcosms, which were fixed with either PFA, DSP (x3) or ethanol after 4 (T4) or 6 hrs (T6) of incubation, and stored for 12 days before click reactions and imaging. Panels a and b have data points (black-filled circles) from the first replicate of each treatment, whereby each data point represents detection ratios calculated from each field of view. For BONCAT comparison image analyses, an average of 3,406 ± 1,533 cells (DAPI stained) were evaluated for each treatment. **c**, Relative abundance bar plots of microbiota community compositions based on 16S rRNA gene amplicon sequencing, presented at the genus level and showing the top 20 most abundant genera. Results from all BONCAT and random sorts are from T6, and results from the microcosm are from T0. Boxplots show the summary statistics where boxes indicate the interquartile ranges (IQR), whiskers show the range of values within 1.5x IQR, and the main horizontal lines indicate the medians.

We next sorted BONCAT-positive cells from glucose incubations (T6) and from “random” cells (bulk sorts of all detectable events) (T6) by FACS (50,000 events per sample) (**Fig. S4**), reversed fixation, extracted DNA, and performed 16S rRNA gene amplicon sequencing from both DSP- and ethanol-fixed sorts. Comparisons of BONCAT-positive cells versus randomly-sorted cells were performed to identify taxa that were enriched in sorted fractions, and thus, stimulated by glucose ^8,30,32,34^. For both fixation approaches, genes for 16S rRNA were readily PCR amplifiable from sorted cells. No significant differences in alpha diversity were detected between random sorts from ethanol versus DSP fixed fractions (**Fig. S5a**). Beta-diversity analysis showed that fixation explained 23.5% of the variation in microbial community composition among sorted fractions (PERMANOVA, R² = 0.235, p = 0.015), whereas the type of sort (BONCAT-positive versus randomly-sorted cells) explained a larger proportion of the variation (35.2%; PERMANOVA, R² = 0.352, p = 0.002) (**Fig. S5b**). In addition, community structures from DSP- and ethanol-fixed samples were more similar to each other than to the original (T0) unfixed microcosm communities (**Fig. 3c and Fig. S5b**). This indicates that the glucose amendment and the sorting process itself, rather than the fixative, had most influence on the recovered community composition. Comparisons of taxonomic relative abundances of DSP- or ethanol-fixed cells from random sorts showed Gram-negative taxa were collectively 13% lower and Gram-positive taxa were collectively 13% higher in ethanol-fixed random sorts compared to DSP-fixed random sorts, although not significantly different (**Fig. S5c**). This is likely due to the higher propensity for Gram-negative bacteria to lyse during fixation and storage in ethanol solutions ^35,36^, and is consistent with the single cell mass loss in ethanol-fixed samples (**Fig. 2a**). Examination of the presence-absence of ASVs from DSP- or ethanol-fixed cells from random sorts showed that a high proportion of ASVs were detected in both sorts, with 324 overlapping, and 35 and 20 ASVs unique to ethanol- and DSP-fixed cells, respectively (**Fig. S5d**). This indicates that DSP fixation does not limit the recovery of ASVs from sorted cells relative to ethanol-fixed cells.

To identify taxa that had significantly higher relative abundances in BONCAT-positive sorts versus random sorts, for cells fixed with either DSP or ethanol, we performed statistical comparisons of 16S rRNA gene relative abundances of genera from these cell sorts. This identified seven and three genera as significantly higher in BONCAT-positive sorts versus random sorts for ethanol- and DSP-fixed cells, respectively (**Table S2**). The genus *Parabacteroides* was determined as significantly higher in BONCAT-positive sorts from both fixation treatments. This aligns with earlier findings that enhanced carbohydrate availability promotes *Parabacteroides* proliferation ^37^. The major discrepancy was for the genus *Bacteroides*. Relative abundance of *Bacteroides* was reduced in random sorts for ethanol-fixed samples compared to random sorts from DSP-fixed samples (**Fig. 3c and Table S2**). Together, these results suggest a potential loss of Gram-negative cells (primarily *Bacteroides*) after ethanol fixation, and DSP as a more reliable fixative for detecting enrichment of BONCAT-positive taxa in comparison to controls.

## DSP enables targeted mini-metagenomics and single-cell genomics

We next tested if DSP fixation enables enrichment of target taxa and improved genome recovery by FISH-FACS and mini-metagenomics. For this, we fixed a human fecal sample with DSP (x2), sorted 60,000 cells of targeted taxa by FISH-FACS targeting *Agathobacter* (EREC996/EREC1252 ^38^), *Faecalibacterium* (FPRAU644 ^39^), *Phocaeicola* (formerly *Bacteroides*) (BVULG1016/BADO374 ^40,41^), and *Lachnospirales* (EREC482 ^42^), and sequenced the mini-metagenomes of sorted cells (**Table S1**). We also sequenced the metagenome of the original fecal sample. Metagenomic read mapping revealed 2–55-fold enrichment of the intended target populations in the mini-metagenomes compared to the original fecal sample (**Fig. S6; Table 1 and S3**). The sorted FPRAU644-stained cells showed strong enrichment of *Faecalibacterium duncaniae* (16-fold) and *Gemmiger* species (12–20-fold), which are targeted by the probe (**Table 1 and S3**). Use of the *Lachnospirales*-targeted EREC482 probe for FISH-FACS enriched diverse species of this order, including *A. rectalis*, *A. faecis, Anthropogastromicrobium aceti*, and *Coprococcus aceti* (8–17-fold) (**Table 1 and S3**). Furthermore, of the FISH-targeted taxa, medium- to high-quality MAGs were recovered from all sorted samples (**Table S3**) ^43^, providing further support that DSP fixation preserves the cellular and genomic integrity of microbial cells. It further shows that DSP fixation enables FISH-FACS mini-metagenomics-based recovery of high-quality genomes of target cell populations.

**Table 1.** Enrichment of target bacterial taxa in mini-metagenomes derived from DSP-fixed human fecal sample by FISH–FACS. The enrichment factor of a target taxon is calculated as the ratio of the percentages of metagenome reads from the unsorted sample and the sorted cell fraction that were mapped to a selected reference genome. An enrichment factor of >1 and <1 indicates fold enrichment and depletion, respectively. FPRAU644, BVULG1016/BADO374, EREC996/EREC1252, and EREC482 correspond to enrichment factors of target taxon genomes in specific probe-defined and sorted cell fractions. Selected reference genomes of probe-targeted taxa were used for read mapping. **Table S3** is an extended version of Table 1 including genome statistics for MAGs that were recovered after FISH-FACS. Grey shading highlights genomes targeted by the respective probe sets.

| Genomes of reference organisms targeted by the applied FISH probes (Genbank accession number) | Enrichment factor in mini-metagenomes [% mapped reads (FISH-FACS)/% mapped reads (total metagenome)]<br>Probes used for FISH-FACS |  |  |  |
| --- | --- | --- | --- | --- |
|  | FPRAU644 | BVULG1016 + BADO374 | EREC996 + EREC1252 | EREC482 |
| *Faecalibacterium duncaniae* (GCF_010509575.1) | 16.0 | 0.3 | 1.8 | 0.9 |
| *Gemmiger quicibialis* (GCF_027689445.1) | 20.3 | 0.7 | 5.7 | 0.7 |
| *Gemmiger formicilis* (GCF_900167555.1) | 12.6 | 1.3 | 2.5 | 1.4 |
| *Phocaeicola vulgatus* ATCC 8482 (GCF_000012825.1) | 0.0 | 7.1 | 0.0 | 0.0 |
| *Phocaeicola dorei* (GCF_013009555.1) | 0.0 | 2.3 | 0.0 | 0.0 |
| *Agathobacter rectalis* (GCF_022453685.1) | 0.7 | 0.7 | 55.0 | 9.4 |
| *Agathobacter faecis* (GCF_001405615.1) | 0.3 | 0.4 | 3.1 | 5.7 |
| *Roseburia hominis* A2-183 (GCF_000225345.1) | 0.3 | 0.4 | 0.4 | 3.7 |
| *Roseburia lenta* (GCF_014287435.1) | 0.0 | 0.1 | 0.2 | 1.4 |
| *Anthropogastromicrobium acetii* (GCF_020687335.1) | 1.5 | 0.2 | 1.3 | 16.9 |
| *Coprococcus acetii* (GCF_025567285.1) | 0.4 | 1.0 | 1.3 | 8.5 |

To determine whether DSP fixation is compatible with single-cell genome amplification, we first sorted individual cells from three in-house cultured isolates, *Escherichia coli* EDL933 (Gram-negative), *Agathobacter rectalis* (Gram-positive), and *Methanobrevibacter smithii* (Archaea), which were fixed with either PFA, DSP (x3), ethanol, or left unfixed. For each isolate, 20 single cells per fixation method were collected via FACS into a 96-well plate and amplified using MDA. Calculation of the average crossing point (Cp, i.e., the cycle at which the inflection point of MDA occurs) showed that DSP fixed cells amplify at the same time as unfixed cells, indicating that DSP fixation does not hamper DNA amplification of single cells **(Fig. 4**). DSP fixation further outperformed PFA fixation across all isolates, while performing equal or better than ethanol fixation for *A. rectalis* and *E. coli*, respectively (**Fig. 4a,c**). Only for *M. smithii* did ethanol fixed cells amplify significantly faster than all other treatments (**Fig. 4b**), possibly because ethanol could disrupt the pseudomurein cell wall more effectively than PFA or DSP ^44^. To determine the coverage of the amplified genomes, we sequenced unfixed, DSP-fixed, and ethanol-fixed single cells of *E. coli*. Mapping of reads to the *E. coli* genome showed that SAGs of unfixed cells had the highest genome coverage (98.7 ± 1.0%), followed by DSP-fixed (78.8 ± 11.6%), with both significantly higher than ethanol-fixed cells (15.8 ± 7.2%, *P* < 0.001, **Fig. 4d**). All SAGs from DSP-fixed cells were >64% complete and reached up to 99.7% genome coverage (**Table S4**), highlighting the ability to recover medium- to high-quality SAGs from DSP-fixed single isolate cells^43^.

**Figure 4.**
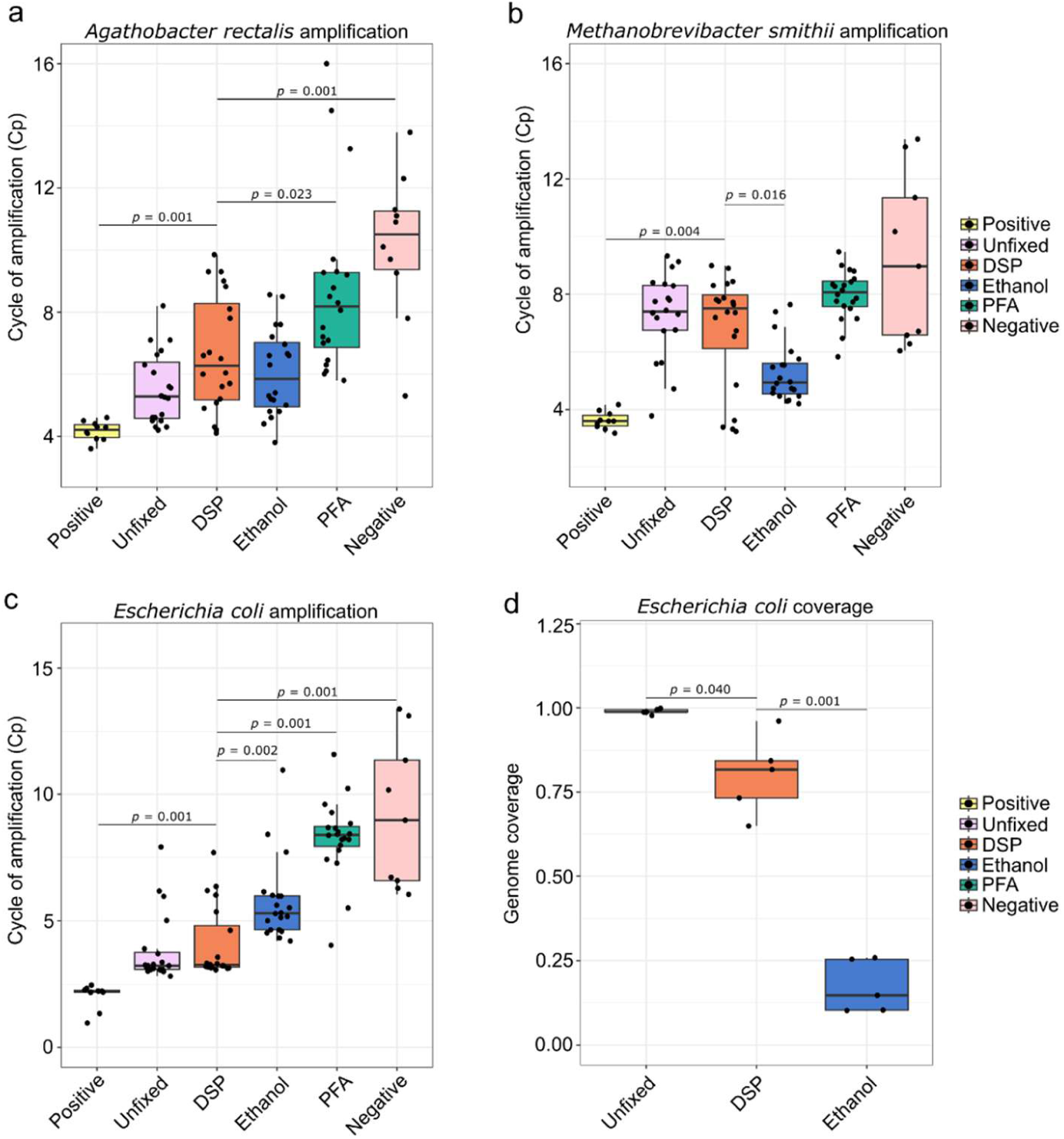
Single amplified genomes (SAGs) are readily obtained from DSP-fixed microbial cells. Average crossing point (Cp, inflection point) values for the real-time MDA amplification curves for *A. rectalis* (**a**), *M. smithii* (**b**), and *E. coli* (**c**). Treatments are colour coded. n = 20 per fixation method (1 cell per well), n = 10 for positive (100 unfixed cells per well) and negative (no cells) controls. **d**, Genome coverage for *E. coli* single cells that were unfixed, DSP fixed, or ethanol fixed (n = 5 per treatment). Only significance between DSP and other treatments is displayed.

To demonstrate that SAGs could be recovered from populations of interest from complex microbial communities, we fixed a human fecal sample with DSP (x3), performed FISH-FACS with the genus-level probe targeting *Phocaeicola* (formely *Bacteroides*) (BVULG1016, ^40^ **Table S1**), and sorted 30 single cells of the target population. The genome of each cell was amplified and sequenced. Metagenomic read mapping showed that 23 of the 30 sorted events (77%) contained *P. vulgatus*, representing a 77-fold enrichment over the unsorted fecal sample (*P. vulgatus* relative abundance ∼1%). Of the 30 sorted events, 11 were pure *P. vulgatus* singlets (single cells in single droplets), from which SAGs with an average completeness of 81 ± 18.4% and contamination of 2 ± 2.5% were recovered (**Table S5a**). The ability to recover high quality SAGs enabled the direct linkage of mobile genetic elements and viruses associated with *P. vulgatus*. GeNomad identified a total of 139 plasmid sequences (plasmid score ≥ 0.7, FDR < 0.05, ≥ 1 plasmid hallmark gene) across the 11 SAGs (**Table S5b**), which clustered into 37 unique plasmid clusters (≥ 90% ANI, ≥ 90% AF). After removal of potential bacterial chromosomal sequences, 23 plasmid clusters remained (**Table S5c**). Eight of these clusters were novel (IMG/PR: pident < 90%, query coverage < 80%) and nine carried mobility genes (**Table S5b,c**).

*P. vulgatus* strains have been reported to carry up to six plasmids, including a megaplasmid and the cryptic plasmid pBI143 (2.7kb) ^45–47^. Here, fragments of pBI143 version 2 (1kb) were identified in 10 out of the 11 SAGs (NODE_725_length_999_cov_10888.828273, **Table S5c)**. GeNomad further identified viral contigs, including 14 prophages, belonging to Monodnaviria (Inoviridae and Microviridae) and Duplodnaviria (Crassvirales and Straboviridae, FDR < 0.05, ≥ 1 viral hallmark gene), with some classified as high-quality or complete by CheckV ^48^ (**Table S5d**). While Crassvirales and Strabovirida have been computationally predicted to infect *P. vulgatus* via CRISPR spacer matching ^49,50^, direct evidence has been absent for all identified viral groups. Together, these findings demonstrate that DSP-based FISH-FACS enables the recovery of high-quality SAGs from complex communities, and allowing the direct identification of host-virus and host-plasmid interactions.

## Conclusion

Here, we present DSP as a reversible fixative for microbial cells that preserves cellular integrity for fluorescence imaging while enabling downstream DNA amplification and sequencing. DSP fixation preserved microbial cell mass for at least several months, was compatible with FISH with fluorescence signals comparable to standard PFA fixation, and supported CARD-FISH, GenomeFISH, and BONCAT, demonstrating broad applicability across phylogeny- and activity-based fluorescence labelling approaches. DSP performed consistently across samples, taxa, and cell types, with experiments independently conducted by different researchers at two institutes, demonstrating robustness across laboratory settings. Finally, FISH-FACS of DSP-fixed human fecal samples enabled up to 77-fold enrichment of targeted taxa and recovery of high-quality MAGs and SAGs.

Combining fluorescence-based cell identification with whole-genome sequencing can provide higher-resolution insights into microbial ecology and genomic potential. DSP fixation can facilitate the identification of viruses and plasmids associated with individual cells or defined populations and enable recovery of high-quality genomes from rare community members that are difficult to capture by conventional metagenomics. Its preservation of nucleic acid integrity, cellular structure, and buoyant mass also opens possibilities for targeted single-cell transcriptomics and proteomics ^51,52^. Combining DSP fixation with cell isolation approaches other than FACS could further broaden its applications ^53^. For example, laser-capture microdissection could link molecular information to the spatial organization of microbial populations within biofilms and host tissues ^54,55^. Taken together, DSP provides a versatile link between fluorescence-based identification and genome-resolved analysis of microbial cells, with broad potential across environmental, clinical, and host-associated microbiome research.

## Materials and Methods

### Fixation comparison experiments with pure cultures

#### Isolate growth conditions and fixation

*Escherichia coli* and *Bacillus subtilis* DSM 10 were grown in LB medium (10 g L^-1^ tryptone, 5 g L^-1^ yeast extract and 10 g L^-1^ sodium chloride, pH 7) overnight (16 hrs) at 37 °C. *Bacteroides thetaiotaomicron* VPI5482 was grown anaerobically (85% N2, 10% CO2, 5% H2, Coy Labs, USA) in BHI medium (CM1135B, Oxoid) for 16 hrs at 37 °C. 0.25 mL of each culture was pelleted at 14,000 *g* for 5 mins and supernatant was removed. All cells were subsequently washed with 1x phosphate buffered saline (PBS), and fixed in either PFA, ethanol, or DSP. All fixation treatments were performed in triplicate.

For PFA fixation, cells were resuspended in 2% PFA in 1x PBS, and incubated for 4 hrs at room temperature. Cells were then washed twice in 1x PBS, resuspended in 1:1 (v/v) ethanol:1x PBS, and stored at -20 °C.

For ethanol fixation, cells were resuspended and incubated in 1:1 (v/v) ethanol:1x PBS at room temperature for 30 mins, and stored at -20 °C.

For DSP fixation, cells were resuspended in 720 µl of 1x PBS, then 30 µl DSP (TCI Chemicals, Japan) (25 mM in DMSO) was added to make 1 mM final concentration, and cells were mixed by inversion of tubes three times. Cells were incubated for 10 mins at room temperature, with mixing by inversion every 5 mins. For multiple treatments with DSP (DSP x2 and x3) cells were pelleted and resuspended in 720 µl of 1x PBS and 30 ul of DSP (25 mM in DMSO). After DSP fixation, Tris-HCl (pH 7) was added to 5 mM final concentration and incubated for 5 mins at room temperature to quench the DSP. Cells were then washed twice with 1x PBS, resuspended in 1:1 (v/v) ethanol:1x PBS, and stored at -20 °C.

#### FISH, microscopy, image processing, and data analysis

FISH experiments were performed after 1.5, 2, and 3.5 months of cell storage at -20 °C. Each replicate of cells processed with each fixation method was subjected to FISH in triplicates. For this, fixed cells were pipetted onto 10-well microscope slides (Marienfeld), then air-dried at 46 °C. The slides were dipped in 1x PBS for 5 s to reduce salts, then air-dried with a stream of compressed air. Cells were hybridized in standard CARD-FISH hybridisation buffer ^56^ with 25% formamide and probe EUB338-I (FITC-labelled) (**Table S1**) at a final probe concentration of 0.25 µmol. Slides were incubated for 2 hrs at 46 °C and then washed for 10 mins at 48 °C in standard FISH wash buffer appropriate for hybridisations with 25% formamide ^24^. Slides were dipped in ice- cold dH2O for 5 s to remove salts, then air-dried with compressed air. Cells were counterstained with 4,6-diamidino-2-phenylindole (DAPI, 10 µg mL^-1^ in 1x PBS), washed with 1x PBS, and air-dried with compressed air. Cells were mounted with Citifluor (Citifluor, UK) and imaged using an inverted Leica TCS SP8X CLSM (Leica, Germany) using appropriate excitation/emission settings for DAPI and FITC, and a 63x objective. Identical microscopy settings were used for each imaging session, i.e., identical zoom (1.2x), gain settings, and line and frame pass numbers. Each well was imaged a minimum of 3-4 times in a random area of the well. Images were analysed using DAIME software (v 2.1) ^57^. For this, images of *E. coli* and *B*. *thetaiotaomicron* cells were subject to biomass detection by “Edge Detection” with Dark threshold “15”, and relative fluorescence values were obtained using the formamide series evaluation tool. For statistical comparisons of relative fluorescence, Kruskall-Wallice tests with Bonferroni corrections and followed by post-hoc Dunn’s test were performed. Box plots were generated from the treatment average values using R.

### Fixation comparison experiments with environmental samples and CARD-FISH

#### River water sample collection and fixation

River Water was collected from the Danube River, Vienna, Austria (48°14’57.15"N 16°23’27.27"E) on March 24 2016, in sterile 500 mL Schott bottles and transported to the laboratory the same day for fixation. 10 mL of water were vacuum filtered onto polycarbonate filters (GTTP type, 0.2-µm pore size) (Millipore, cat. no. GTTP02500). For PFA fixation, filters were incubated in 4% PFA in 1x PBS for 2 hrs at room temperature, and then washed twice by incubating in fresh 1x PBS for 2 mins. Filters were air-dried at 46 °C and stored at -20 °C. For DSP fixation, filters were incubated three times in fresh 1 mM DSP in 1x PBS for 10 mins, and then washed twice by incubating in fresh 1x PBS for 2 mins. Filters were air-dried at 46 °C and stored at -20 °C. For ethanol fixation, filters were incubated in 1:1 (v/v) ethanol:1x PBS at room temperature for 30 mins, air-dried at 46 °C, and stored at -20 °C. Each fixation treatment was performed in triplicate.

#### CARD-FISH, microscopy, image processing, and data analysis

CARD-FISH hybridisations were performed after 10 days of storage at -20 °C. For CARD- FISH, whole filters were dipped cell-side down in pre-dissolved 0.1% low-melt agarose, dried on parafilm, and removed using 100% ethanol and air-dried. Endogenous peroxidases were inactivated by incubating filters in 0.1% H2O2 (Sigma) in 1x PBS for 2 mins at room temperature. Filters were washed twice by incubating in 50 mL of fresh 1x PBS for 5 mins each. Permeabilization was performed by covering filters in lysozyme (10 mg mL^-1^; Sigma-Aldrich) for 15 mins at room temperature. Filters were washed twice by incubating in 50 mL of fresh 1x PBS for 5 mins each. Filters were then cut into small sections and incubated at 46 °C for 3 hrs in CARD- FISH hybridisation buffer with 20% formamide for Horseradish Peroxidase (HRP)-labelled probes EUB-I-II-III mix, ALF968 and NONEUB, and 35% formamide for GAM42a plus unlabelled competitor probe BET42a (**Table S1**). Probes were added at 1 µl of probe (50 ng µl^-1^ stock) per 300 µl of CARD-FISH hybridisation buffer. Each hybridisation was performed in triplicate. Filters were washed twice by incubating in 50 mL of fresh 1x PBS for 10 mins each. Tyramide signal amplification was performed using standard protocol for 15 mins at 46 °C and using Alexa-488 labelled tyramides ^56^. Filters were washed twice by incubating in 50 mL of fresh 1x PBS for 10 mins each. Filters were counterstained with DAPI (10 µg mL^-1^ in 1x PBS) for 5 mins, washed with 1x PBS for 5 mins and then again in fresh 1x PBS for 30 mins, then air-dried. Filters were mounted with Citifluor (Citifluor, UK) and imaged using an inverted Leica TCS SP8X CLSM (Leica, Germany) using appropriate excitation/emission settings for DAPI and FITC, and using a 63x objective. The same microscopy settings were used for imaging each filter, i.e., identical zoom (1.2x), gain settings, and line and frame pass numbers. Each filter was imaged a minimum of 3- 4 times in a random area of the filter. Images were analysed using DAIME software (v 2.1) ^57^. For DAIME processing of images of DAPI stained cells, custom thresholding was used with Low = 200 and High = 255, and ignore objects <2 and >55 pixels. For DAIME processing of images of CARD-FISH-stained cells (green fluorescence), custom thresholding was used with Low = 50 and High = 255, and ignore objects <2 and >55 pixels. Object counts were obtained using the formamide series evaluation tool in DAIME. Statistical comparisons were performed as described above for evaluation of FISH on pure cultures.

#### Marine sediment sample collection, and fixation

Shallow (*c*. 2-5 cm below surface) estuarine marine sediment (Aveiro, Portugal) (N 40° 33’ 41.503’’; W 8° 45’ 22.649’’) were collected at low tide on July 27 2019, and added to sterile 50 mL Schott bottle. Sediment was filled to the very top, sealed with a lid and electrical tape to exclude oxygen where possible, and transported to the laboratory for processing within a week. All following fixation treatments were performed in triplicate.

For PFA fixation, 0.5 mL of sediment was collected into 2 mL tubes, and 1.5 mL of 4% PFA in 1x PBS was added and mixed by vortexing for 5 s. Samples were fixed for 4 hrs at room temperature, washed twice with 1x PBS and resuspended to 1.7 mL with 1x PBS, before proceeding with Nycodenz cell clean-up method described below.

For DSP fixation, 0.5 mL of sediment was collected into 2 mL tubes and resuspended in 1 mL of 1x PBS. Then 60 µl of DSP (25 mM stock in DMSO) was added, samples were vortexed for 5 s, and incubated at room temperature for 10 mins with vortexing every 5 mins. This was repeated two more times. Samples were resuspended to 1.7 mL in 1x PBS to proceed with Nycodenz cell clean-up.

For ethanol fixation, 0.5 mL of sediment was suspended up to 2 mL of 1:1 (v/v) ethanol:1x PBS and incubated at room temperature for 30 mins. Samples were washed twice with 1x PBS and resuspended to 1.7 mL with 1x PBS to proceed with Nycodenz cell clean-up.

#### Cell separation, CARD-FISH, and microscopy for marine sediment samples

For cell clean-up from marine sediments, cells were first dislodged from particles using chemical and sonication treatments ^58^. For chemical treatment, sodium pyrophosphate was added to 0.1% final, and Tween 20 to 0.5% final, and tubes were shaken horizontally for 30 mins at 700 rpm at room temperature (‘speed’ 5) using a Vortex Genie 2 vortexer (Scientific Industries). Samples were sonicated with 25% power for 30 s using setting ‘5’, while on ice (UW 2070 needle, Bandelin Electronics). Samples were then brought to room temperature to allow Nycodenz to be added, and brought up to 4 mL with 1x PBS in 13.2-ml Thinwall Polypropylene Tubes (Beckman Coulter). 2 mL of Nycodenz solution (80% w/v) (Alere Technologies, cat. no. 1002424) was carefully injected under the sample suspension. Samples were centrifuged for 90 min at 4 °C in an SW 41 Ti Swinging-Bucket Rotor (Beckman Coulter) at 14,000 *g*, with no deceleration when stopping. The total supernatant including the Nycodenz fraction but not sediment particles was carefully collected to new 15 ml tubes. For DSP fixation treatments, DSP was also added to the collected cells at 1 mM final, vortexed for 3 s, and incubated at room temperature for 10 mins with vortexing every 5 mins. Finally, ethanol was added to each replicate of each treatment to make- up to 50% (v/v), and samples were stored at -20 °C.

CARD-FISH and DAPI staining was performed after 3 days of storage at -20 °C with HRP- labelled EUB-I-II-III mix as described above for river water CARD-FISH experiments, except 200 µl of cleaned-up cells stored in 1:1 (v/v) ethanol:1x PBS were vacuum filtered to polycarbonate filters (GTTP type, 0.2-µm pore size) (Millipore, cat. no. GTTP02500), and approximately 5 mL of 1x PBS was washed through the filter by vacuum after. Microscopic imaging, image analyses and statistical comparisons of sediment cells was performed as described above for river water CARD-FISH experiments with the following differences: for DAIME processing of images of CARD-FISH stained cells, custom thresholding was used with Low = 100 and High = 255, and ignore objects <2 pixels.

### GenomeFISH

#### Sample collection and fixation

A fecal sample was collected from one healthy donor under the Queensland University of Technology ethics agreement (Approval Number 5203) and transferred into an anaerobic chamber within 15 mins. Approximately 50 mg of sample was resuspended in 1000 μL of filter sterilised (Millex-GP 0.22 μm PES) anaerobic diluent and vortexed for 10 s before being filtered through a 35 μm FACS cell strainer (Falcon; 352235) into a sterile 1.5 mL microfuge tube. Filtered faecal biomass was fixed in DSP as described above for river water and stored in 1:1 (v/v) ethanol:1x PBS at -20°C.

#### GenomeFISH

To visualise *Agathobacter rectalis* in the fecal sample, GenomeFISH probes were generated from an *A. rectalis* strain (AHMB0000188) obtained from the Australian Human Microbiome Biobank (AHMB). Briefly, DNA was extracted from 1 mL overnight cultures using the DNeasy PowerBiofilm Kit (Qiagen, Germany), following the manufacturer’s instructions. DNA was fragmented and fluorescently labelled with Alexa546 for GenomeFISH on the fecal sample as outlined previously *^7^*. In brief, fixed biomass was applied to polytetrafluoroethylene (PTFE) printed slides and dried at 37°C for 10 mins. Cells were dehydrated in an ethanol series (50%, 80%, 100%; 3 mins each) and permeabilized with lysozyme (10 mg mL^−1^ lysozyme in 1x PBS, 0.05 M EDTA, and 0.1 M Tris–HCl, pH 8) for 1 h at 37°C. Slides were washed in MilliQ and 100% ethanol, and further permeabilized with hydrochloric acid (0.2 M, 15 mins at room temperature). To each well, 10 μl of hybridization buffer was added (35% deionized formamide, 5X SSC (750 mM NaCl, 0.075 mM sodium citrate, pH 7), 20% dextran sulfate, 20 mM EDTA, 0.25 mg mL^−1^ sheared salmon sperm DNA, 0.25 mg mL^−1^ yeast RNA, 1x Denhardt’s blocking solution, and 0.1% SDS), together with 1 μl of genome-targeted probe (to a final concentration of 2.5 ng μl^−1^). Labelled calf thymus DNA probes were used as a negative control. Slides were incubated for 40 mins at 85°C to denature the target DNA, followed by hybridization overnight at 46°C. After hybridization, slides were submerged in pre-warmed wash buffer (80 mM NaCl, 100 mM Tris–HCl pH 8, 5 mM EDTA, and 0.001% SDS) for 15 mins at 48°C, followed by a 20 mins wash at room temperature in 1x PBS, 1 min in MilliQ, and 2 mins in 100% ethanol. Cells were counterstained with DAPI (1 μg ml^−1^) for 15 mins at room temperature and washed in MilliQ and 100% ethanol. Slides were mounted with VECTASHIELD PLUS antifade mounting medium (VECTOR Laboratories) and microscopic analysis was performed with a Stellaris5 (Leica) laser scanning confocal microscope.

### Single-cell buoyant mass measurements of unfixed and fixed human fecal samples

#### Sample collection and fixation

To characterise the effect of storage time on microbial single-cell buoyant mass, fecal samples were obtained from two individuals (A and B). Fecal samples were collected from another ten healthy adults (balanced sex, 25-42 years, varied diets) for the direct comparison of single- cell buoyant mass between DSP and PFA fixation. Each of the ten donors provided samples at three time points (n = 30) six days to two months apart. Each time, donors submitted triplicate fecal subsamples from the most recent stool (1-4 h after defecation). Approximately 0.3–0.4 g of fecal samples from each biological triplicate was pooled to obtain ∼1 g of homogenized fecal material. Samples were resuspended in 30 mL ice-cold 1x PBS (pH 7.4), and rehydrated on ice for 10 mins. After thorough homogenization by shaking and vortexing for 25–30 mins with intermittent cooling, large fecal fragments were manually disrupted with a spatula. To remove debris while retaining microbial cells in suspension, the slurry was centrifuged at low speed (1000 rpm, ∼110 x g) for 2 mins at 4 °C. This procedure was repeated three times, with transfer of the supernatant after each spin. The aliquots of pooled supernatants with microbial cells were preserved with different fixatives. For PFA fixation, samples were resuspended in 3% PFA and incubated for 1 h at room temperature. Cells were pelleted, washed with ice-cold 1x PBS, resuspended in 1x PBS, and stored at 4 °C. For DSP (x2) fixation, freshly prepared 25 mM of DSP in DMSO was added to 0.5 mL sample to a final concentration of 5 mM, followed by incubation for 10 mins at room temperature. After centrifugation, cell pellets were resuspended in 1 mM DSP, incubated for a further 10 mins, washed once in ice-cold 1x PBS, resuspended in 1x PBS, and stored at 4 °C. For ethanol fixation, samples were mixed 1:1 with 99% molecular-grade ethanol (final concentration 50%), incubated for ∼1.5 h at room temperature, washed twice with ice-cold 1x PBS, resuspended in 1x PBS, and stored at 4 °C. 500 µL of each fixed sample was stored in 1x PBS at 4°C.

#### Single-cell buoyant mass measurements

Single-cell buoyant mass was measured using a LifeScale-Research Instrument (Affinity Biosensors), a commercial implementation of the suspended microchannel resonator principle. Briefly, particles or cells transit through a microfluidic channel embedded within a vibrating cantilever, which induces a transient shift in the resonant frequency that is proportional to the buoyant mass of the particle ^59–61^. Buoyant mass reflects the particle volume and the density difference between the particle and the surrounding fluid. Single-cell buoyant mass measurements have previously been established for biological particles and cells with femtogram- scale sensitivity ^27,62^.

Here, the instrument was configured for high-throughput single-cell detection, with a nominal lower buoyant-mass detection limit of approximately 14 fg under the optimized flow conditions. Cells were aspirated directly from sample wells and passed through the cantilever sensor. Particle detection, event calling and real-time conversion of resonance-frequency shifts to buoyant mass were performed using the instrument’s built-in software with user-defined flow- rate and sensitivity settings (load duration 1.5 s, maximum well duration 3 mins, clear inlet every 20 s, sensor wash after each 16 samples). Instrument calibration was verified using polystyrene beads with National Institute of Standards and Technology-certified diameter and density values (Thermo Scientific 4000 Series Monosized Particles; 1.0, 1.6, 2.0, 3.0 and 4.0 µm, and occasionally 5.0 and 6.0 µm) as done previously ^4^. To minimize non-cellular background particles, all media and buffers were filtered immediately before use through 0.1-µm polyethersulfone syringe filters (Minisart® High Flow, Sartorius; cat. no. 16553-K). Buffer controls were analyzed before sample acquisition to confirm a sufficiently low particle background (<1 × 10^5^ particles mL^−1^), comparable to ultrapure water controls.

The combined effects of fixative and storage time on single-cell buoyant masses were measured in samples from donors A and B. Each sample was left unfixed or fixed with DSP, PFA, or ethanol and stored for one day and 42 days in 1x PBS at 4°C. For Sample A, the numbers of cells analyzed (Day1/Day42) were: DSP 12,253/12,508, PFA 18,006/12,720, EtOH 11,658/5,171, and unfixed 7,514/5,061. For Sample B, the corresponding counts were: DSP 5,381/5,815, PFA 7,681/6,702, EtOH 6,608/4,191, and unfixed 3,422/5,224. Buoyant mass was log10-transformed for visualization and kernel density ridgelines were plotted for each sample and treatment.

Single-cell buoyant masses were measured in DSP- and PFA-fixed samples (n = 30 each) from the study cohort. Median single-cell buoyant masses of corresponding PFA- and DSP-fixed samples were analyzed using (i) Spearman rank correlation (ρ and p-value) and (ii) an ordinary least squares linear regression model DSP ∼ PFA, extracting the regression fit and R². Differences in fixation medians were further evaluated with a paired Wilcoxon signed-rank test (p-value). Across samples, the number of single-cell measurements analyzed per fixation condition ranged from 2,236 to 19,712 for DSP (mean 7,346) and from 2,788 to 46,500 for PFA (mean 10,652).

### Fixation comparison experiments for BONCAT

#### Sample collection, BONCAT and fixation

A fresh fecal sample was collected from a healthy adult individual, transferred into an anaerobic chamber (85% N2, 10% CO2, 5% H2, Coy Labs, USA), and processed within 90 mins of defecation. A 10% (w/v) fecal slurry was prepared by adding 1x PBS to the sample, followed by thorough vortexing for 2 mins. The sample was then left for 10 min to settle larger particles. 2 mL of the pre-settled mixture was transferred to a new 50 mL falcon tube and diluted with sterile 1x PBS to obtain a final 0.5% (w/v) slurry. The mixture was vortexed, and larger particles were allowed to settle. Aliquots (5 mL) of this fecal slurry were subjected to the following treatments, in triplicates for each: A) 500 nM glucose amendment with 50 µM L-azidohomoalanine (AHA); B) 500 nM trehalose amendment with 50 µM AHA; C) no substrate amendment with 50 µM AHA (control for background AHA incorporation). An additional single control with 500 nM trehalose amendment with no AHA (negative control) was performed. These were incubated anaerobically at 37 °C for 4 and 6 h before sampling for fixation.

For PFA fixation, 500 µl of sample was washed twice in 1 volume of ice-cold 1x PBS,resuspended in 4% PFA in 1x PBS, and incubated at 4 °C for 2 h. Samples were then washed twice with 1 mL of 1x PBS, resuspended in 500 µl of 1:1 (v/v) ethanol:1x PBS, and stored at -20 °C.

For ethanol fixation, 500 µl of sample was washed twice in 1 volume of ice-cold 1x PBS, resuspended in 500 µl of 1:1 (v/v) ethanol:1x PBS, and stored at -20 °C.

For DSP fixation, 500 µl of sample was washed twice in 1 volume of ice-cold 1x PBS, resuspended in 400 µl of 1x PBS and 100 μl of DSP (25 mM DSP in DMSO), vortexed for 5 s, and incubated at room temperature for 10 mins. Samples were pelleted, resuspended in 480 µl of 1x PBS and 20 μl of DSP (25 mM DSP in DMSO), vortexed for 5 s, and incubated at room temperature for 10 mins. This was repeated once more. Samples were then washed twice with 1 mL of 1x PBS, resuspended in 500 µl of 1:1 (v/v) ethanol:1x PBS, and stored at -20 °C.

#### Click-chemistry, microscopy, image processing, and data analysis

Click-chemistry reactions of AHA-incubated microcosms (BONCAT labelling experiments) were done after 3 and 12 days of storage at -20 °C. 15 µl of sample were added to wells of 10- well glass microscope slides (Marienfeld), air-dried at 46 °C, dipped in ice-cold dH2O for 5 s to remove salts, then air-dried with a stream of compressed air. A dye pre-mix was prepared by adding 1.25 μl of 20 mM CuSO4 solution, 2.50 μl of 50 mM Tris[(1-hydroxypropyl-1H-1,2,3-triazol- 4-yl)methyl]amine (THPTA) and 0.30 μl of alkyne Cy5 dye (Carboxyrhodamine 110 Alkyne, ClickChemistryTools; 10 mM in DMSO), and allowed to react for 3 mins at room temperature in the dark. Meanwhile, a solution was prepared by mixing 221 ul of 1x PBS, 12.5 μl of 100 mM freshly prepared sodium ascorbate and 12.5 μl of freshly prepared 100 mM aminoguanidine hydrochloride, which was added to the dye pre-mix and mixed once by inversion of the tube. This dye solution (15 μl) was then added to microscope wells to cover samples, added to a 50 mL tube that included tissue paper covered with 200 μl 1x PBS, and incubated for 30 mins at room temperature in the dark.

For microscopy, 10 µl of samples were added to wells of a 10-well glass microscope slide (Marienfeld), air-dried at 46 °C, dipped in ice-cold dH2O for 5 s to remove salts, then air-dried with a stream of compressed air. Cells were counterstained with DAPI (10 µg mL^−1^ in 1x PBS) for 5 mins, washed with 1x PBS, and air-dried with a stream of compressed air. Cells mounted with Citifluor (Citifluor, UK) and imaged using an inverted Leica TCS SP8X CLSM (Leica, Germany) using appropriate excitation/emission settings for DAPI and Cy5, and using a 63× objective. Identical microscopy settings were used for each imaging session, i.e., identical zoom (1.2×), gain settings, and line and frame pass numbers. Each well was imaged 4-5 times in a random area of the well. Images were analysed using DAIME software (v 2.1) ^57^. DAIME processing of images of DAPI stained cells, custom thresholding was used with Low = 50 and High = 255; and for DAIME processing of images of Cy5 BONCAT-labelled cells, custom thresholding was used with Low = 50 and High = 255. Object counts were obtained using the formamide series evaluation tool in DAIME.

#### BONCAT-FACS

For BONCAT-FACS, the click-chemistry reaction was performed in solution, according to Hatzenpichler *et al.* 2015 ^63^. Briefly, 250 μl of fixed samples (T6 glucose incubation samples, stored for 141 days at -20°C) were pelleted (3,500 *g*, 5 mins at room temperature), washed once using 96% ethanol, and resuspended in 221 μl 1x PBS. A dye premix solution was prepared by mixing 1.25 μl of a 20mM CuSO4 solution, 2.50 μl of a 50 mM solution of THPTA (tris[(1- hydroxypropyl-1H-1,2,3-triazol-4-yl)methyl]amine, ClickChemistryTools) and 0.30 μl of alkyne Cy5 dye (Carboxyrhodamine 110 Alkyne, ClickChemistryTools; 10 mM in DMSO). The dye mix was let to react at room temperature in the dark for 3 mins. In the meantime, 12.5 μl of a 100 mM solution of sodium ascorbate and 100 mM solution of aminoguanidine hydrochloride (both from Carl Roth GmbH; freshly prepared) were added to the sample. Dye premix was then added, samples were inverted once, and incubated for 30 mins at room temperature in the dark. Afterwards, samples were washed 3 times by centrifugation (3500 *g*, 5 mins room temperature) and resuspended in 1x PBS. All samples were transferred into flow cytometry tubes and passed through a 35 µm pore cell strainer cap (Corning) to remove larger particles or cell aggregates.

Samples were analyzed on a BD FACSMelody Cell Sorter (BDBiosciences). Background signals from the instrument and PBS were identified using the operational parameters forward scatter (FSC) and side scatter (SSC). Fluorescence events (BONCAT, Cy5 dye) were detected using a 561 nm laser and a 697/58 nm filter. Gating for BONCAT-positive cells was determined based on samples for which a click reaction was performed without dye. Instrument parameters and gating strategy were kept constant during acquisition of all samples. For each sample, 50,000 cell events were sorted in purity mode.

#### DNA isolation and 16S rRNA gene sequencing from BONCAT-sorted fractions

Sorted cell fractions (30-50 μl) were collected into 1.5 mL tubes and immediately stored at -80°C until further processing. For DNA isolation, DSP-fixed cells were thawed on ice and DSP crosslinking was reversed by adding 1 M DTT to the tube containing the DSP-fixed cells suspended in 1x PBS to a final concentration of 20 μM of DTT. Tubes were incubated at 37°C for 30 mins with occasional mixing. To enhance cell lysis, 150 μL of enzymatic lysis buffer (20 mM of TRIS-HCl, pH 8.0, 2 mM of Na-EDTA, 1.2% v/v of TritonX-100, 20 mg mL^−1^ of Lysozyme from chicken egg purchased from Sigma Aldrich) was added to both ethanol-fixed and DSP fixation- reversed sorted cells. Samples were incubated for 60 mins at 37°C with gentle shaking. The DNA was then extracted using the DNeasy Blood and Tissue kit (QIAGEN), according to the manufacturers instructions. Nucleic acids were eluted by adding 25 μl of nuclease-free water (Carl Roth GmbH), incubated at room temperature for 5 mins and centrifuged for 1 min. Eluent was stored at -20°C until further usage. Extraction of the kit buffer only and of 1x PBS collected from the cell sorter (sorting gated on electronic noise) were performed in parallel with the samples, as controls.

16S rRNA gene amplicon sequencing was performed by the Joint Microbiome Facility (JMF) of the University of Vienna and the Medical University of Vienna (project number JMF- 1908-3). Bacterial 16S rRNA genes were amplified from DNA samples using a two-step PCR barcoding approach^64^ with primers targeting most bacteria (S-D-Bact-0341-b-S-17, 5′- CCTACGGGNGGCWGCAG-3′ and S-D-Bact-0785-a-A-21, 5′-GACTACHVGGGTATCTAATCC-3′). First-step PCRs were performed in triplicate (12.5 μl vol per reaction) with the following conditions: 1x DreamTaq Buffer (Thermo Fisher), 2 mM MgCl2 (Thermo Fisher), 0.2 mM dNTP mix (Thermo Fisher), 0.2 µM of forward and reverse primer each, 0.08 mg ml^-1^ Bovine Serum Albumin (Thermo Fisher), 0.02 U Dream Taq Polymerase (Thermo Fisher), and 0.5 µl of DNA template. Conditions for thermal cycling were: 95°C for 3 mins, followed by 30 cycles of 30 s at 95°C, 30 s at 55°C and 60 s at 72°C, and finally 10 mins at 72°C. Triplicates were combined for barcoding (8 PCR cycles). Barcoded samples were purified and normalized over a SequalPrep™ Normalization Plate Kit (Invitrogen) using a Biomek® NXP Span-8 pipetting robot (Beckman Coulter), and pooled and concentrated on innuPREP PCRpure Kit columns (Analytik Jena). Indexed sequencing libraries were prepared with the Illumina TruSeq Nano Kit as described previously ^64^, and sequenced in paired-end mode (2x 300 bp; v3 chemistry) on an Illumina MiSeq following the manufacturer’s instructions. The workflow systematically included four negative controls (PCR blanks, i.e., PCR-grade water as template) for each 90 samples sequenced.

#### Analysis of 16S rRNA gene amplicon sequences

Amplicon pools were extracted from the raw sequencing data using the FASTQ workflow in BaseSpace (Illumina) with default parameters. Input data was filtered for PhiX contamination with BBDuk (BBTools, Bushnell B, sourceforge.net/projects/bbmap). Demultiplexing was performed with the python package demultiplex (github.com/jfjlaros/demultiplex) allowing one mismatch for barcodes and two mismatches for linkers and primers. DADA2 ^65^ was used for demultiplexing amplicon sequencing variants (ASVs) using a previously described standard protocol ^65^. FASTQ reads 1 and 2 were trimmed at 230 with allowed expected errors of 4 and 6, respectively. Taxonomy was assigned to 16S rRNA gene sequences based on SILVA taxonomy (release 138) using the naïve Bayesian classification method as implemented in DADA2 ^65^.

Sequencing of kit buffer extraction controls yielded a low number of reads (2 and 11 reads) assigned to ASV_qdi_vob (*Pseudomonas*). Sequencing of FACS-sorted PBS controls (three replicates, gated on electronic noise) yielded 14, 20, and 80 reads, respectively. These reads were predominantly assigned to ASV_qdi_vob (*Pseudomonas*) and ASV_qee_9fh (*Alcaligenes*), with a smaller number of reads classified as other *Pseudomonas* or Gammaproteobacteria ASVs. As these ASVs dominated controls and were present at very low levels in the sorted cell fractions, they were considered contaminants and excluded from subsequent analyses. After removal of contaminant ASV reads, amplicon sequencing yielded an average of 13,148±4,699 reads per sample belonging to a total of 474 ASVs. Good’s coverage was >98.5% for all samples. ASVs with fewer than 10 total reads across all samples were removed ^66^. Pruned amplicon sequence libraries were analysed using the vegan (v2.4.3) and phyloseq (v1.30.0) packages of the software R (https://www.r-project.org/, R 3.4.0). To evaluate the effects of fixation on Gram-positive versus Gram-negative bacteria in BONCAT-associated experiments, 16S rRNA gene amplicon sequences were assigned to either type based on their phylum-level classifications, i.e., Actinobacteria and Firmicutes were assigned as Gram-positives, and Bacteroidetes, Proteobacteria and Lentisphaerae were assigned as Gram-negatives. Other sequences not from these above listed phyla belonged to “unclassified” sequences and were on average 0.3% relative abundance across all samples, and a maximum of 0.7%. No archaea were detected.

### Recovery of mini-metagenomes from human faeces by FISH-FACS

#### Sample collection, fixation, and FISH

A freshly collected human fecal sample (0.5 g) was resuspended in 1 ml 1x PBS and washed twice by centrifugation at 5,000 × g to remove debris and soluble material. The washed pellet was resuspended in 1 ml 1x PBS and treated twice with 5 mM DSP for 15 mins at room temperature, with a 1x PBS wash between treatments. After DSP removal, samples were stored at −20 °C in a 20% (v/v) glycerol:1x PBS buffer.

To minimize cell loss and reaggregation during FISH, 10–20 µl subsamples were transferred to microcentrifugal filter units with 0.2 µm polytetrafluoroethylene (PTFE) membranes (Thermo Fisher). The glycerol-PBS buffer was removed, and cells were washed with 1x PBS by centrifugation at 2,000 x g in a mini-centrifuge (Benchmark Scientific) until no supernatant remained. For FISH, samples were washed and pre-equilibrated 200 µl hybridization buffer (without probe) and centrifuged in a mini tabletop centrifuge. Subsequently, 100 µL of hybridization buffer containing Cy3-labeled oligonucleotide probes was added (see next paragraph, and Table S1 for details). The filter units were placed in fresh 1.5 mL microcentrifuge tubes, sealed, and incubated for 3 h at 46 °C. After hybridization, samples in the microcentrifugal filter units were washed twice by centrifugation for 30 s at 2,000 × g in a pre-warmed (48 °C) rotor of a standard centrifuge (Eppendorf 5804R). To this end, the pelleted, hybridized stool subsamples were resuspended and washed twice in 500 µL of pre-warmed washing buffer (48 °C), with incubation for 10–15 mins each. The samples were then pelleted again, resuspended in ice-cold 1x PBS buffer, transferred to sterile 2 ml tubes and ultrasonicated for 15 s using a Bandelin Sonopuls (cycle 2, 25% power; Berlin, Germany) to disaggregate cells. Samples were kept on ice until FACS analysis.

Fluorescence *in situ* hybridization (FISH) was performed using a set of group- and species-specific 16S rRNA-targeted oligonucleotide probes with double-labeled Cy3 fluorophores (DOPE-FISH) ^67^. The following probes were applied: an equimolar mix of BVULG1016 and BADO374 for *Phocaeicola vulgatus* and *dorei*, FPRAU644 for *Faecalibacterium prausnitzii*, an equimolar mix of EREC996, EREC1252, and competitors for *Agathobacter rectalis*, and EREC482 for most members of the order *Lachnospirales*, including *Agathobacter rectalis* (**Table S1**). The nonsense probe NON338 and a no probe control served as negative controls to assess nonspecific binding and background fluorescence.

#### Flow cytometric sorting

To enhance discrimination from background debris, all samples were stained with SYBR Green I (1× final concentration; Thermo Fisher). Flow cytometric analysis and sorting were performed using a FACSMelody (BD Biosciences) equipped with 488 nm and 561 nm lasers for detection of green (SYBR Green, 527/32 nm) and red (Cy3, 582/15 nm) fluorescence, respectively. Data acquisition and analysis were conducted with FACSChorus software. A low side scatter (SSC) threshold was applied to exclude electronic noise and small noncellular debris from the buffer background. Events displaying Cy3 fluorescence equal to or greater than the NON338 control were gated as probe-positive. For each target population, at least 60,000 double- gated (SYBR Green- and Cy3-positive) cells were sorted and collected for DNA extraction.

To assess sorting success and purity, approximately 1,000 cells per target population were directly sorted onto a small area (∼1 mm²) of a 0.2 µm polycarbonate membrane, stained with DAPI, and examined using an epifluorescence microscope (Zeiss Axio Imager M1 equipped with an AxioCam MRm). For each probe, 90–95% of DAPI-stained cells displayed overlapping FISH signals, confirming high sorting specificity.

Prior to DNA extraction, sorted cells were centrifuged at 5,000 × g for 15 mins, and DSP cross-linking was reversed using DTT as described above for BONCAT sorted cells. The original DSP-fixed, unsorted stool sample served as a control to evaluate target population enrichment following FACS. For metagenomic sequencing, 0.2 mL of the original stool sample was processed for reversal of DSP fixation and DNA extraction.

#### DNA extraction and sequencing

Metagenome sequencing was performed by the JMF (project number JMF-2411-04). DNA was extracted using the QIAamp DNA Mini Kit (Qiagen, Germany), incorporating a lysozyme pre- treatment step to enhance cell lysis and optimize DNA recovery ^68^. Individual samples were prepared for sequencing using the NEBNext® Ultra™ II FS DNA Library Prep Kit for Illumina according to the manufacturer’s instructions. The fragment sizes and quality of the individually prepared libraries was checked on an Agilent High Sensitivity D1000 Tape, using an Agilent Tape Station 4150 device. Following fluorometric DNA quantification with Qubit 4.0 ™, the barcoded libraries were equimolarly pooled. The final insert size and concentration of the pool was determined (Agilent High Sensitivity D1000 Tape and Qubit™) and sequenced on an Illumina NovaSeq 6000 (2x 100 bp, SP flowcell).

### Quality control of reads

A quality control snakemake v8.28.3 workflow was used to trim the read libraries based on quality scores. SAMtools v1.20’s ^69^ bam2fq was used to convert bam files to fastq sequences for each read library. The read libraries were quality checked using fastQC v0.12.1 (https://github.com/s-andrews/FastQC) and quality statistics were merged using multiQC v1.21^70^. Adapters were trimmed and phiX contamination was removed using BBDuk (part of BBMap v39.06)^71^ to a minimum of 50 base pairs in length. Reads were k-trimmed from the right with a kmer of 21, minimum kmer of 11 and hamming distance of two along with the “tpe” and “tbo” options. Quality trimming was performed from the right with a Q-score of 15. The quality-controlled read libraries were quality checked using fastQC v0.12.1 and quality statistics were merged using multiQC v1.21 for comparison to the non-trimmed quality. BBMap’s reformat.sh was used to interleave the read libraries.

#### Metagenomic assembly and binning

The read libraries were assembled using SPAdes v4.0.0 ^72^ with kmers of 21 to 121 in steps of 10 and the “meta” mode. Contigs under 1000 base pairs were removed from the resulting assembly using reformat.sh of BBMap (https://github.com/bbushnell/BBTools). MetaBAT v2.15 ^73^ was used to generate putative MAGs without coverage information using a minimum length of 1500 base pairs (minimum possible length). Coverage information for the assembly was generated with BBMap with a minimum identity of 98% using the read libraries. The resulting bam files were sorted using SAMtools v1.20 ^74^. A depth file was generated using jgi_summarize_bam_contig_depths (from MetaBAT2) and the assemblies were binned into putative MAGs again using the depth file from MetaBAT2. All putative MAGs were dereplicated using dREP v3.5.0 ^75^. Additional dereplication was performed with selected genomes from NCBI. The taxonomy of the dereplicated MAGs was determined using GTDB-tk v2.4.0’s classify_wf ^76^ with database r220, skipping ANI screening. Relative abundance was calculated using the “genome” workflow of CoverM v0.7.0 ^77^.

### Single cell genomics

#### Isolates growth conditions and fixation

The ability to recover single amplified genomes from DSP fixed cells was first tested on three in-house cultured isolates, including a Gram-negative (*Escherichia coli* EDL933), Gram-positive (*Agathobacter rectalis*, AHMB0000188), and Archaea (*Methanobrevibacter smithii* PS).

*E. coli* was grown as described above, *A. rectalis* was grown in anaerobic modified TY medium at 37°C, and *M. smithii* in BRN-RF10 media with 150kPa of H2/CO2 (80:20) incubated with 120 rpm rotational shaking at 37°C.

For each isolate, 1 mL of cells were collected from cultures at late exponential stage, pelleted, and washed twice in 1x PBS. Cells were either fixed in PFA, ethanol, DSP, or kept unfixed. For PFA and ethanol fixation, cells were fixed overnight at 4°C in 4% PFA (w/v) or 1:1 (v/v) ethanol:1x PBS, respectively. Cells were fixed in DSP as described above for the fecal samples. Unfixed and fixed cells were stored at -80 °C in 1x PBS with 20% glycerol (v/v).

#### Fecal sample collection and in-solution FISH

A fecal sample was collected from one healthy donor under the Queensland University of Technology ethics agreement (Approval Number 5203) and processed as described above for GenomeFISH visualisation. Fixed faecal sample (150 µl) was pelleted at 14,000 g for 10 mins and cells were permeabilized with 50 µl lysozyme (0.05 mg mL^-1^ lysozyme in 1x PBS, 0.05 M EDTA, and 0.1 M Tris–HCl, pH 8) for 30 min at 37 °C. Cells were pelleted, washed in 1x PBS, and dehydrated for 3 mins each in 50, 80, and 96% (v/v) ethanol. Cells were pelleted again, the ethanol removed, and the cell pellet air dried. Cells were subsequently hybridized overnight at 46°C in solution with 10 µl hybridization buffer (900 mM NaCl, 20 mM Tris-HCl, 1 mM EDTA, 30% formamide, and 0.01% SDS) and 1 µl of the S-S-Bvulg-1017-a-A-21 (double labelled in Alexa488), targeting *Phocaeicola* (*Bacteroides*) (**Table S1**). The NONEUB probe^19^ was used as a negative hybridization control. After hybridization, samples were transferred into a centrifuge with a rotor preheated at 46 °C and centrifuged at 14,000 g for 15 mins at 40 °C to minimize unspecific probe binding. Samples were incubated in pre-heated wash buffer for 15 mins at 48 °C, centrifuged for 15 mins at 14,000 g to remove buffer, and washed in ice-cold 1x PBS. Cells were finally centrifuged at 14,000 g for 10 mins at 4 °C and resuspended in 1 mL of ice-cold 1x PBS for single cell sorting.

#### Single cell sorting and amplification

Single cells were sorted with a FACS Aria Fusion Flow Cytometer (BD Bioscences). Briefly, isolate samples stored in 20% glycerol were defrosted on ice, diluted 1:1000 in 0.2 µm filtered 1x PBS, filtered through a 40 µm FACS cell strainer (Falcon; 352235), and individually sorted into a 384 well plate. The sort window for isolates was determined based on forward (FSC) and side (SSC) scatter plots from the small particle detector. Cells processed through in-solution FISH were sonicated for 20 seconds (20% power, Omni Sonic Ruptor 4000), filtered through a 40 µm FACS cell strainer (Falcon; 352235), and individually sorted into a 96 well plate. To sort FISH positives, the bacterial community was first identified using FSC and SSC plots with thresholds set on 0.2 µm filtered 1x PBS. Events that were positive for DAPI (405 - 450/50) and S-S-Bvulg- 1017-a-A-21 (in A488 - 488 530/30) were subsequently collected from this region of scatter where bacteria typically appear. Negative controls of the faecal sample labelled with the NON338 probe were also run to ensure no false positive events. Doublets and chains were excluded using Area vs Height and Width vs Height. Prior to each sort, the instrument was sterilized by soaking the entire fluidics system in 1% sodium hypochlorite for a minimum of 30 mins and UV crosslinking the sheath fluid (sterile 1x PBS) for 1 h.

Multiple displacement amplification (MDA) of single cells was performed as in Engelberts et al. ^7^ and the inflection crossing point (i.e. the point of exponential amplification before the reaction reaches a plateau) measured. Single amplified genomes (SAGs) from wells with successful amplification were purified using the ZYMO DNA Clean & Concentrator Kit (ZYMO Research), following the manufacturer’s instructions and DNA concentrations were measured with the Qubit dsDNA BR Assay Kit (Invitrogen).

#### DNA extraction from fecal sample

To gain insight into the microbial community of the original fecal samples, total DNA was extracted from the homogenized, unfiltered fecal samples at Microba Life Sciences (Brisbane) using the DNeasy 96 PowerSoil Pro QIAcube HT kit and the QIAcube HT DNA extraction system, following the manufacturer’s instructions.

#### Sequencing

DNA from the fecal sample and the best amplified MDA products of unfixed, ethanol fixed, and DSP fixed *E. coli* single cells (n = 5 per treatment) was sent for sequencing to Microba Life Sciences (Brisbane). DNA from *P. vulgatus* SAGs (n = 30) was sent for sequencing to the Centre for Microbiome Research (CMR, Brisbane). At Microba Life Sciences, libraries were prepared using the Illumina DNA Prep Library Preparation Kit as per the manufacturer’s instructions, with unique dual indexes and PhiX spike in at 2%. Samples were sequenced on the NovaSeq6000 in 2 × 150 bp format. At CMR, libraries were prepared using the QIAseq FX DNA Library UDI Kit (Cat. No. 180479, QIAGEN), following the manufacturer’s instructions. Samples were sequenced on an Illumina NovaSeq X system using standard XLEAP-SBS chemistry according to the manufacturer’s recommended workflow. DNA of all SAGs was sequenced to 0.5 Gbp and DNA of the fecal sample was sequenced to 5 Gbp.

#### Data analysis

Reads from the fecal sample and the *E. coli* and *P. vulgatus* SAG were quality filtered using RastQC v0.1.0 ^78^. The genome coverage of the *E. coli* and *P. vulgatus* SAGs were subsequently calculated by mapping reads to the reference genomes of *E. coli* and *P. vulgatus* using CoverM v0.7.0 ^77^ with the ‘genome’ command. To eliminate biases caused by varying sequencing depth, the *E. coli* sequences were randomly subsampled to an even sequencing depth prior to read mapping.

Reads of the fecal sample and *P. vulgatus* SAGs were further assembled and binned using Aviary v0.13.0 ^79^. In short, reads were assembled using SPAdes v4.2.0 ^72^ and binned with the binning algorithms MetaBAT2 v2 ^73^, MaxBin v2 ^80^, CONCOCT ^81^, VAMB ^82^, SemiBin v2 ^83^, and Rosella ^84^. The best representative metagenome assembled genome (MAG) produced from the different binning algorithms was selected using DASTool v1.1.7 ^85^. CheckM2 ^86^ was used to determine completeness and contamination of both the assemblies (for SAGs, i.e. pure singles) and recovered MAGs. Taxonomy was assigned to the SAGs and MAGs with GTDB-Tk v2 ^87^ using GTDB database R226.

Putative plasmid and viral sequences were identified in the *P. vulgatus* SAGs using GeNomad v1.11.0 ^88^. Only sequences with a plasmid score of at least 0.7 and false-discovery rate smaller than 0.05, while encoding at least one geNomad plasmid hallmark, were kept. Plasmids were subsequently dereplicated based on Reciprocal Blast Hits, using 90% ANI and 90% Aligned Fraction (AF), with the largest sequence kept per cluster. Reference sequences of each cluster were mapped using Blastn v2.10.1 (output format 6) against two databases. To identify potential chromosomal contamination, sequences were mapped to microbial MAGs recovered from the original fecal sample and classed as contaminants if they showed >90% identity and >80% query coverage. To assess novelty, sequences were mapped to IMG/PR (downloaded 26 June 2026) and classed as novel if they showed <90% identity and <80% query coverage. For viruses, only sequences with a score of at least 0.7 and false-discovery rate smaller than 0.05, while encoding at least one geNomad viral hallmark, were kept. CheckV v1.1.1 ^48^ was used to determine the quality of the recovered viral contigs.

All statistical analyses were performed in R v4.1.2^6^. As the variance across groups (unfixed, DSP fixed, and ethanol fixed *E. coli* cells) was unequal, *P* values were calculated using the Welch ANOVA, with a Games-Howell post-hoc test. Graphs were created in R v4.1.2^6^ using ggplot2 v3.3.5 and refined in Inkscape v1.2 (https://inkscape.org/).

### Ethics

Sampling and microbiota analyses of feces from healthy human adults was approved by the University of Vienna Ethics Committee (reference numbers 00161 and 00714) and the Queensland University of Technology ethics agreement (Approval Number 5203). All subjects gave their informed consent for inclusion before they participated in the study.

## Acknowledgements

Isolates for the single cell work (i.e., *A. rectalis*, AHMB0000188) were sourced from the Australian Human Microbiome Biobank operated by Queensland University of Technology (https://ahmb.com.au/). We also acknowledge the technical support of the Flow Cytometry Facility at the Queensland Brain Institute (QBI) at the University of Queensland (UQ), and especially thank Virginia Nink and Allison S. McInnes for their technical assistance. We thank Ben Roller and Martin Polz for support with single cell buoyant mass measurements, Christian Winter for support with initial FACS analyses, and Jasmin Schwarz, Gudrun Kohl and Julia Ramesmayer from the JMF for support with sequencing. Computational results of this work have been achieved using the Life Science Compute Cluster (LiSC) of the University of Vienna.

## Data availability

The 16S rRNA gene sequences were deposited in the NCBI Sequence Read Archive (SRA) as BioProject Accession PRJNA1537448. Raw sequence data associated from the stool metagenomes before and after FISH-FACS were deposited in the Sequence read Archive (SRA) under project accession PRJNA1537342, corresponding MAGs are available from Zenodo (doi: 10.5281/zenodo.23086984). Raw sequence data, MAGs, and SAGs associated with the single cell work were deposited in the European Nucleotide Archive (ENA) under project accessions PRJEB127646 (fecal raw sequences and MAGs) and PRJEB127628 (SAGs).

## Contributions

KW and AL conceived the original idea, which was extended with JPE and SJM. KW performed all standard-FISH and CARD-FISH fixation comparison experiments and analyses. JPE performed GenomeFISH experiments. MM and BTH performed FISH-targeted mini-metagenomics experiments. NS performed single cell buoyant density experiments. FCP and KW performed BONCAT experiments and analyses. JO and BH performed bioinformatic analyses. JPE performed all single cell genomics experiments, with input from CR and AP. JGV cultured in-house isolates and processed samples for sorting. GWT and SJM contributed essential infrastructure. JPE, KW, and AL interpreted the data and wrote the manuscript with help from all authors.

