## Supplementary Information for "Reversible fixation of microbial cells for targeted genomics"

**General recommendations for DSP fixation of microbial cells and samples**

**Supplementary Tables S1-5 legends**

**Supplementary Figures S1-5**

**General recommendations for DSP fixation of microbial cells and samples**

Here, we provide a protocol with general recommendations for DSP fixation of microbial cultures and different environmental sample types.

Prepare a 25 mM stock of DSP (TCI Chemicals) in DMSO (anhydrous) (Sigma-Aldrich) fresh for each fixation experiment (e.g., dissolve 101.1 mg of DSP in 10 ml DMSO, or scale down for small or few samples). The bottle of DSP powder should be brought to room temperature before opening to avoid condensation, because it is moisture sensitive.

If cells are from an organic-rich culture or environment (e.g., fecal matter, wastewater treatment plant), we recommend to wash cells prior to DSP treatment, e.g., with 1x PBS 1-2 times by pelleting and removal of supernatant, in order to reduce extracellular organics that DSP may react with. After washing, resuspend the sample in 1x PBS (pH 7-9) to a desired end concentration of cells, e.g., 400 μl or 800 μl, or other. Add DSP to a final concentration of 5 mM. Vortex the sample for 3 s. Allow the sample to fix for 10 mins at room temperature, with vortexing for 3 s every 5 mins. While fixing a sample in DSP once is generally sufficient for downstream workflows, we occasionally observed improved FISH signals by repeating the fixation an additional 1-2 times. To do this, after the first fixation, wash the sample by pelleting and removing the supernatant, then resuspend in 1x PBS, and repeat the 5 mM DSP fixation step. If precipitates form, DSP can be added to the sample slowly, and/or vortexed to dissolve. We note that if precipitates appeared, they did not affect downstream experiments. After the final fixation, wash the cells with 1x PBS, and resuspend cells in a desired volume of 1x PBS. Cells can be stored in either 1:1 (v/v) ethanol:1x PBS at -20 ^o^C for at least several months, or 20% glycerol:1x PBS at -80 ^o^C for more long-term storage and to possibly protect DNA integrity. For storage in 20% glycerol:1x PBS, mix the glycerol into the sample well and allow it to penetrate into the cells for 5 mins before freezing.

For cells recovered by filtration, cover cells on the filter in 5 mM DSP in 1x PBS for 10 mins at room temperature, wash the filter by incubating in fresh 1x PBS for 2 mins. Repeat DSP fixation an extra 1-2 times if desired. Perform a final wash in 1x PBS for 5 mins. Air-dry the filter (e.g., at 46 ^o^C), and store dry at -20 ^o^C for long-term storage.

For sediment or soil samples, collect a sample equivalent of 0.5 mL of material into a 2 mL tube and resuspend to a total of 1 mL of 1x PBS. Add DSP at 5 mM final concentration, vortex for 5 s, and incubate at room temperature for 10 mins with vortexing every 5 mins. Wash cells with 1x PBS, then repeat the DSP fixation two more times. This can be scaled-up for larger sample volumes. Resuspend the sample in 1x PBS to a desired volume to proceed with cell clean-up, e.g., in 4 mL 1x PBS for a Nycodenz procedure (e.g., see main Methods). After cells are separated from particles, e.g., via Nycodenz treatment, add DSP to the collected cells at 1 mM final, vortex for 3 s, and incubate at room temperature for 10 mins with vortexing for 3 s every 5 mins. Wash cells in 1x PBS, and store in either 1:1 (v/v) ethanol:1x PBS at -20 ^o^C for at least several months, or 20% glycerol:1x PBS at -80 ^o^C.

After FISH procedures and to prepare fixed cells for FACS, cells stored in ethanol:1x PBS should be washed with 1x PBS to remove ethanol. Cells stored in glycerol:1x PBS can be diluted and used directly. Mild sonication for 20 s should be performed to dissociate cell aggregates, before running cells through a cell strainer (40 microns) for FACS.

To reverse fixation for downstream amplification and sequencing from multiple cells, add DTT to DSP-fixed cells suspended in 1x PBS to a final concentration of 20 μM of DTT. Incubate the cells at 37°C for 30 mins with occasional mixing, after which DNA can be extracted. For single cell genomics, multiple displacement amplification (MDA) reactions can be performed using the REPLI-g Single Cell Kit (QIAGEN) as described previously (i.e., with DTT in the lysis buffer) (Rinke 2018). If using other MDA reagents or kits, include DTT in the lysis step in a similar manner to the REPLI-g Single Cell Kit method. To improve cell lysis, a freeze–thaw cycle/s (10 min at −80°C and 10 min at 55°C) should be added after the lysis of cells for 10 min at room temperature.

**Supplementary Table legends**

**Table S1.** rRNA-targeted oligonucleotide probes used for FISH, including probe name, target organism, sequence, and coverage.

**Table S2.** Overview of BONCAT statistics, including microbial relative abundance and enrichment factor across treatments.

**Table S3.** Enrichment of targeted microorganisms in FISH-FACS mini-metagenomes relative to the source fecal sample, and metagenome assembled genome (MAG) statistics for the targeted microorganisms.

**Table S4.** Overview of single amplified genomes (SAGs) recovered from an *E. coli* isolate.

**Table S5.** Overview of single amplified genomes (SAGs) recovered from a fecal sample through FISH-FACS, including A) *P. vulgatus* SAG CheckM2 statistics, B) putative plasmid contigs in *P. vulgatus* SAGs, C) plasmid clusters in *P. vulgatus* SAGs, and D) putative viral contigs in *P. vulgatus* SAGs.

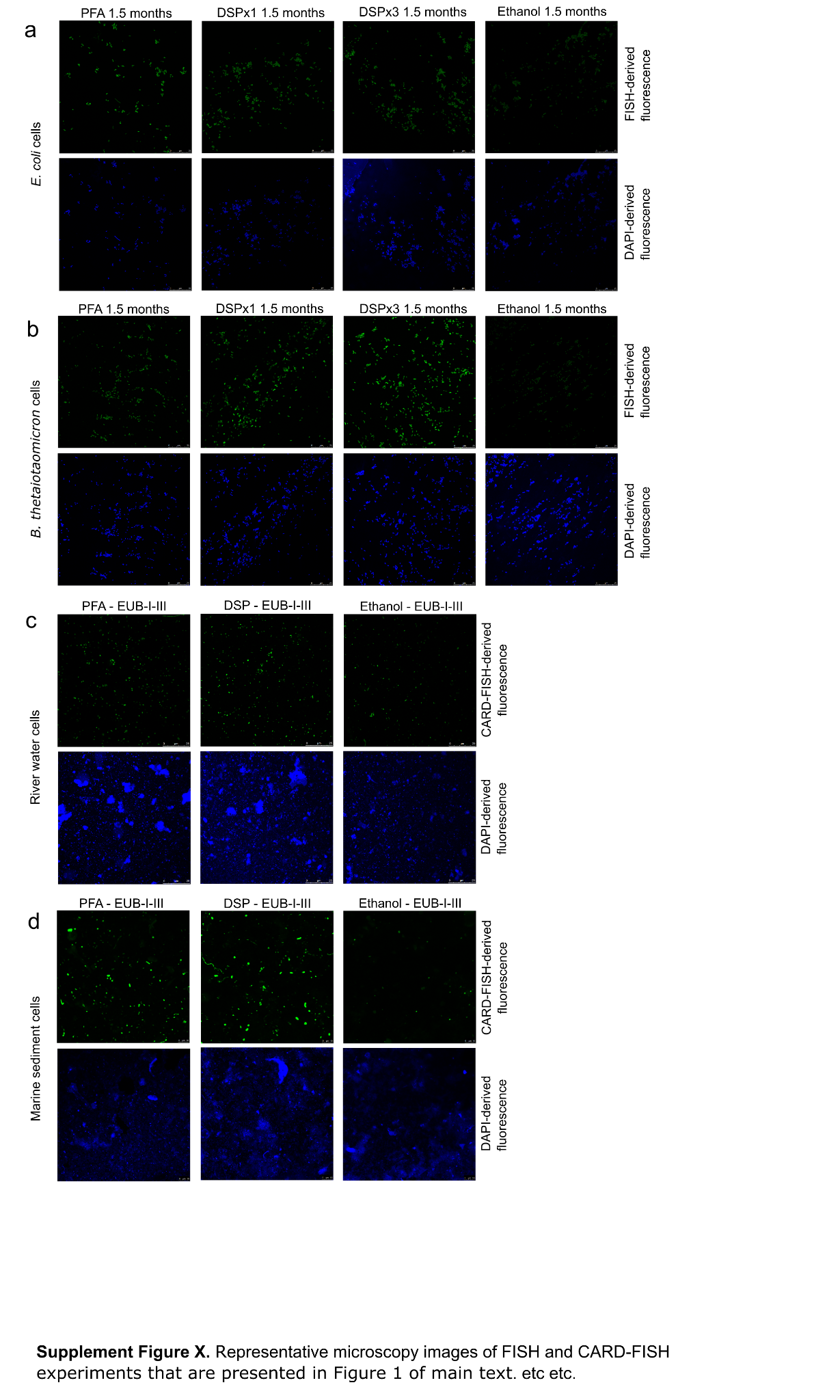

**Figure S1**. Representative microscopy images of FISH and CARD-FISH experiments that are presented in Figure 1 of main text. a, *E. coli* cells from FISH using FITC mono-labelled EUB338-I probe and fixed with either PFA, DSP (x1), DSP (x3), or ethanol. Cells were hybridised and imaged after 1.5, 2, and 3.5 months of storage. **b**, *B. thetaiotaomicron* cells from FISH using FITC mono-labelled EUB338-I probe and fixed with either PFA, DSP (x1), DSP (x3), or ethanol. Cells were hybridised and imaged after 1.5, 2, and 3.5 months of storage. **c**, Cells from river water from CARD-FISH using probes EUB338-I-II-III probe mix targeting most Bacteria. **d**, Cells from marine sediments from CARD-FISH using probes EUB338-I-II-III probe mix targeting most Bacteria. Images with green stained cells are from FISH or CARD-FISH fluorescence, and images with blue stained cells were stained by DAPI. Scale bars in images in panel a-c are 25 µm, and 10 µm in panel d. PFA = paraformaldehyde. DSP = dithio-bis(succinimidyl propionate).

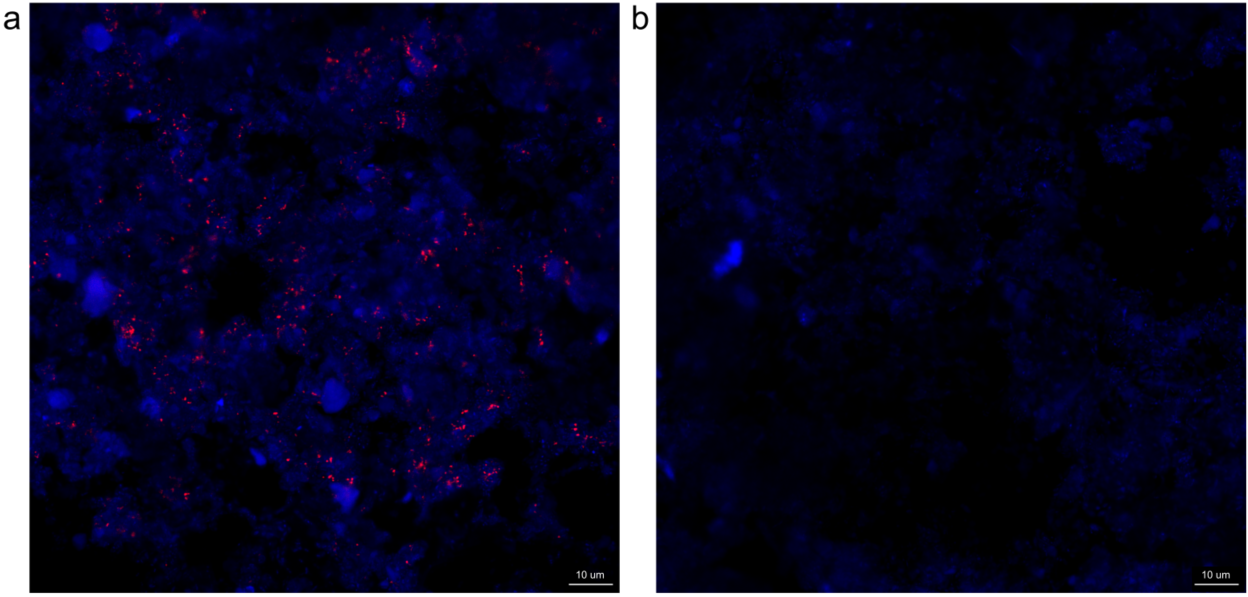

**Figure S2**. GenomeFISH microscopy images of *A. rectalis* in a human fecal sample (**a**) and a negative control (i.e. labelled calf thymus DNA probes, **b**). GenomeFISH signal appears red and DAPI signal blue. Scale bars are 10 µm.

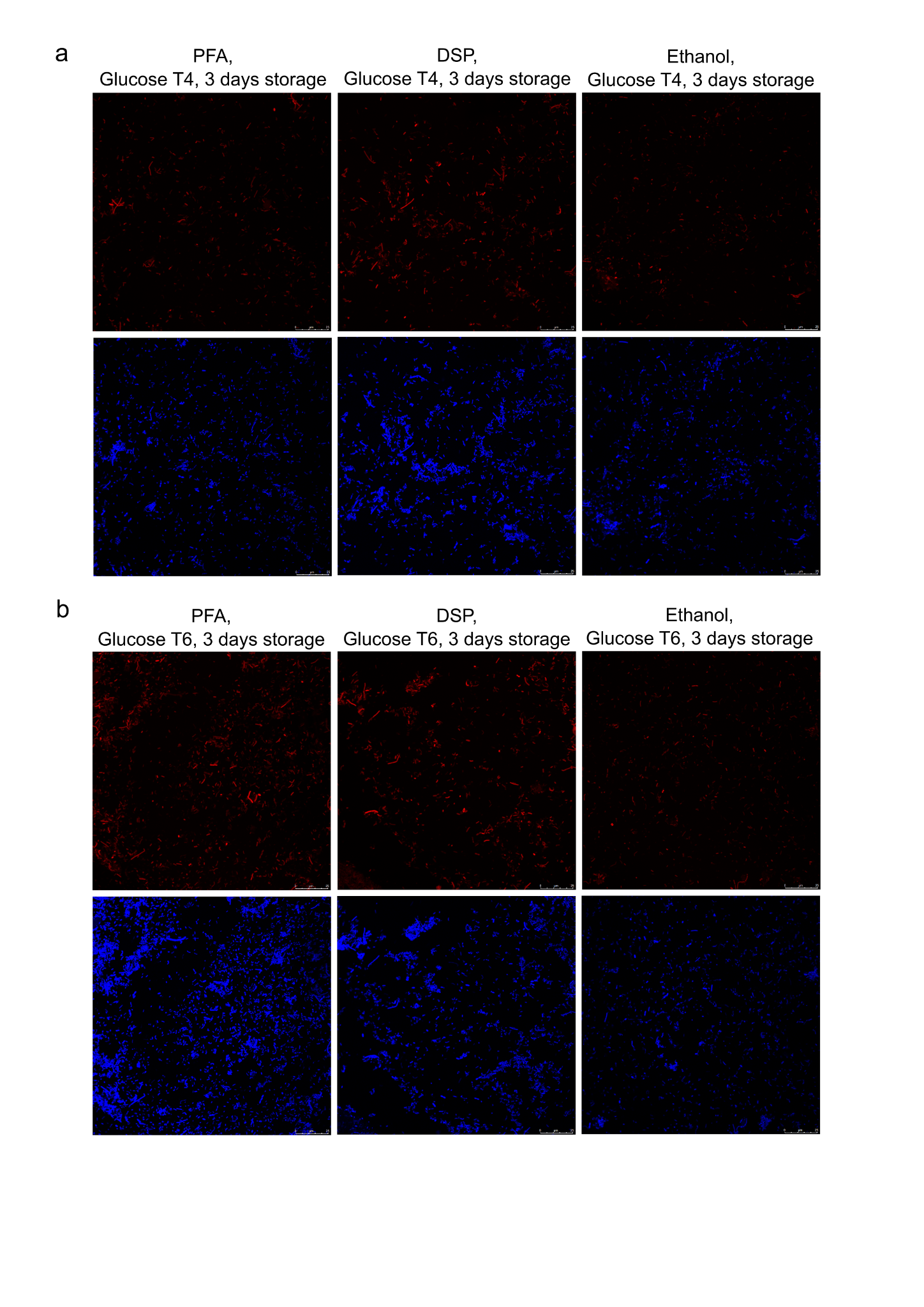

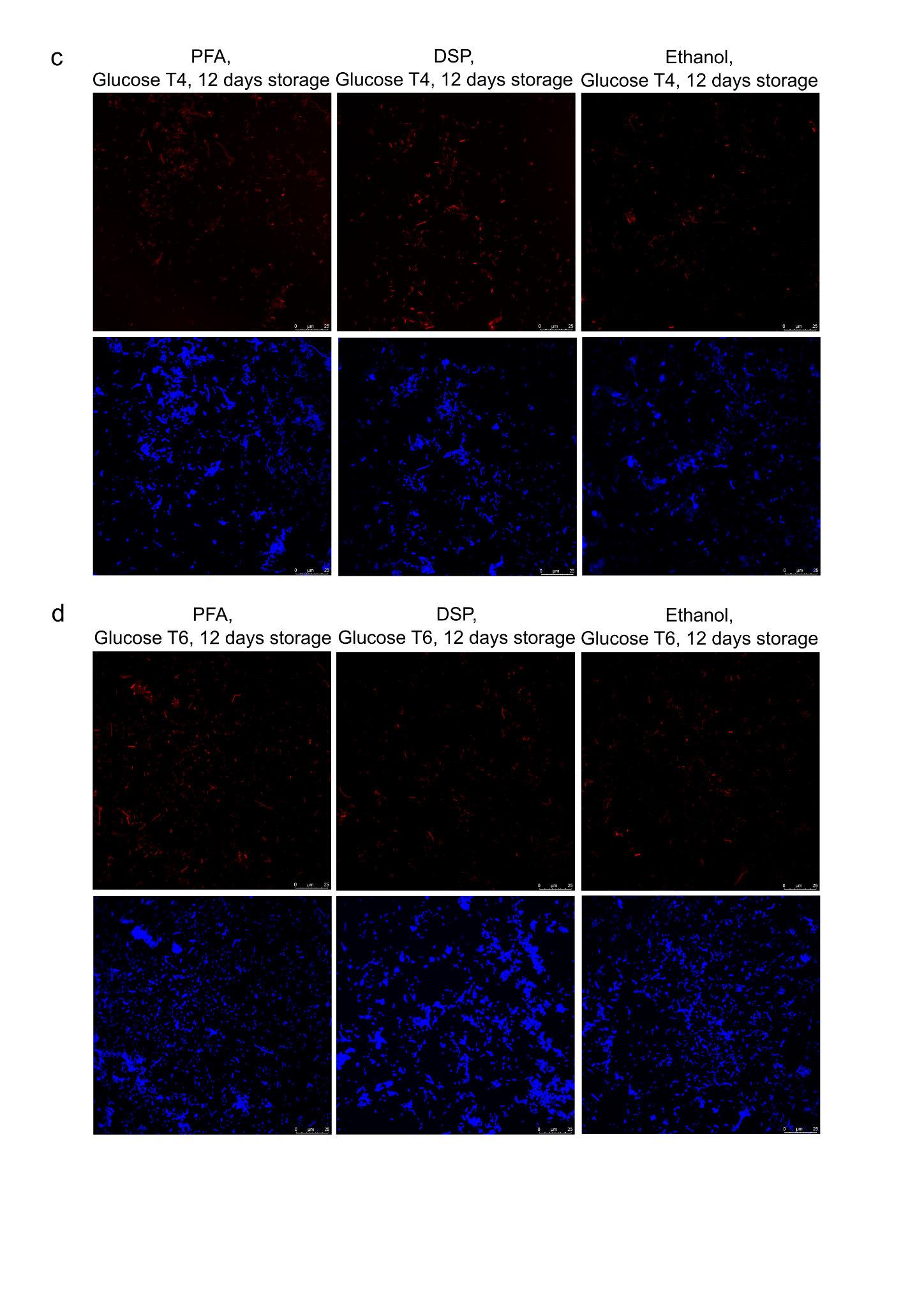

**Figure S3. BONCAT microscopy images.** Representative images of BONCAT experiments of human fecal microcosms incubated with glucose. **a**, Samples were incubated for 4 hours (T4) and BONCAT click reactions were performed after 3 days of storage. **b**, Samples were incubated for 6 hours (T6) and BONCAT click reactions were performed after 3 days of storage. **c**, Samples were incubated for 4 hours (T4) and BONCAT click reactions were performed after 12 days of storage. **d**, Samples were incubated for 6 hours (T6) and BONCAT click reactions were performed after 12 days of storage. Images with red cells are from BONCAT (Cy5) fluorescence, and images with blue stained cells were stained by DAPI. Scale bars are 25 µm.

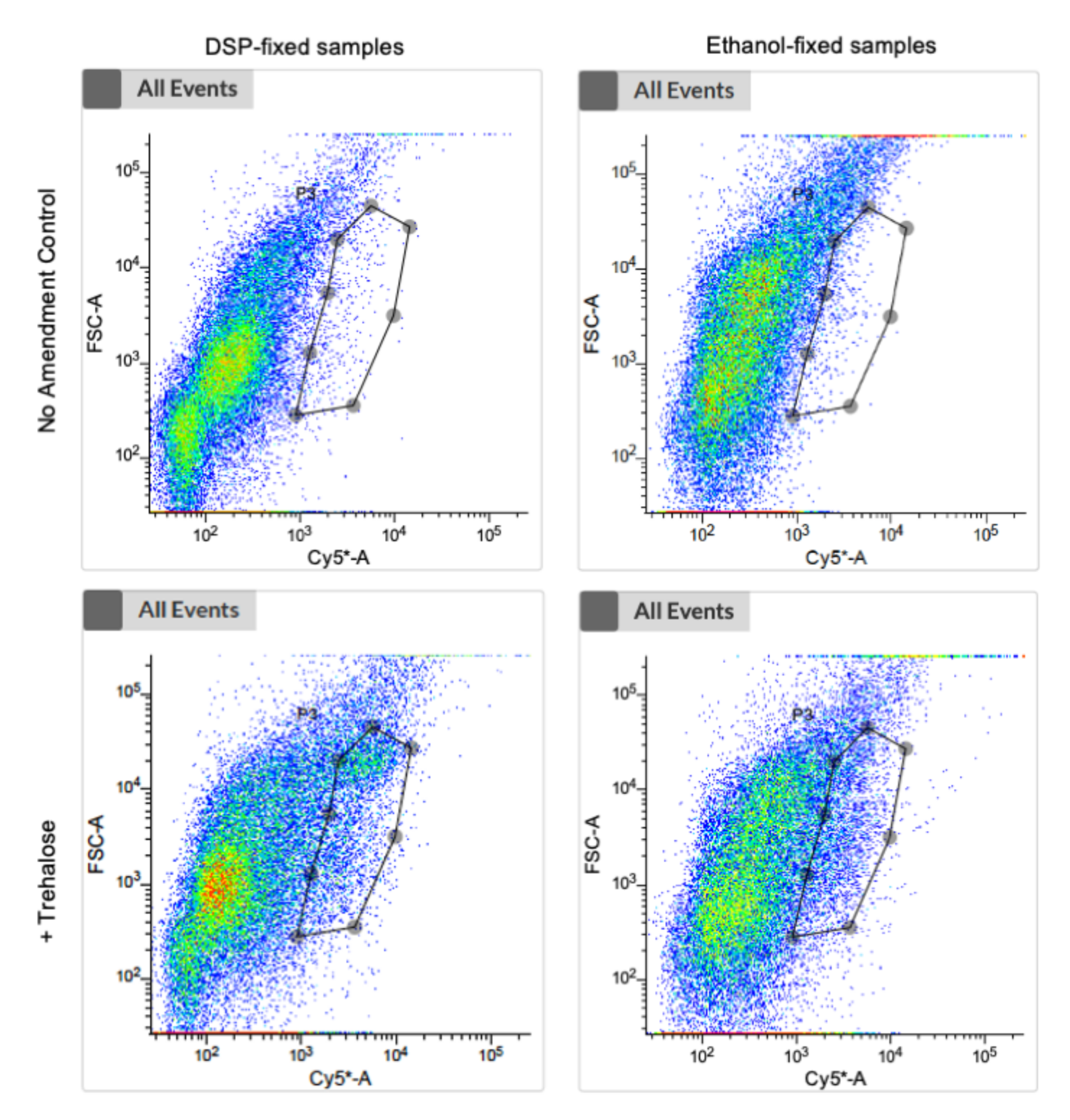

**Figure S4. BONCAT FACS sorting gate images.** Representative scatter plots showing forward scatter (FSC-A) and Cy5-fluorescence (Cy5-A) for samples fixed with either DSP or ethanol, and amended with trehalose or no-amendment controls. Sorting gates for BONCAT-positive cells are shown in each plot (lines connected by grey-filled circles). The same sorting gate was used for sorting BONCAT positive cells from both glucose or trehalose amended samples. For sorting of “random” events, all detected events were sorted.

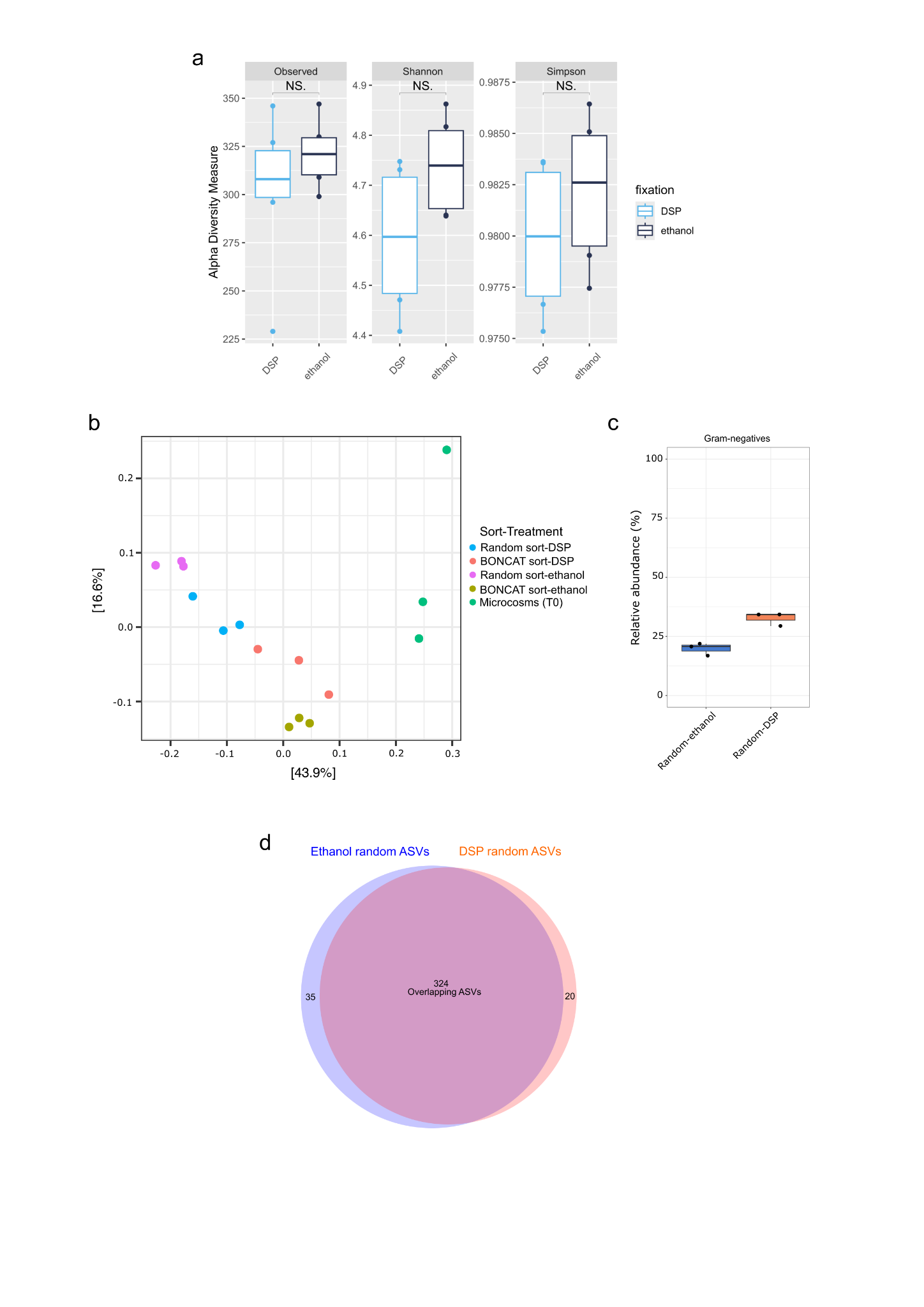

**Figure S5. 16S rRNA gene amplicon supplement analyses.** **a,** Alpha diversity comparisons from random FACS sorts from BONCAT experiments. Alpha diversity measures for Observed ASVs, Shannon and Simpson indexes of 16S rRNA gene diversity for DSP- and ethanol-fixed samples. NS. denotes non-significant difference. Boxplots show the summary statistics where boxes indicate the interquartile ranges (IQR), whiskers show the range of values within 1.5 × IQR, and the main horizontal lines indicate the medians. **b**, Principal Coordinate Analysis (PCoA) of microbiota community compositions based on 16S rRNA gene amplicon sequencing of original microcosm samples (T0, unfixed), BONCAT-positive sorts from ethanol- or DSP-fixed samples from glucose amended microcosms (T6), and random sorts from ethanol- or DSP-fixed samples from glucose amended microcosms (T6), based on a Bray-Curtis dissimilarity matrix. **c**, Relative abundance box plots of microbiota community compositions based on 16S rRNA gene amplicon sequencing with taxa grouped according to the Gram-stain classification of cell wall types for phyla. **d**, Venn diagram showing the numbers of overlapping 16S rRNA gene ASVs detected in random sorts of cells from ethanol- and DSP-fixed samples.

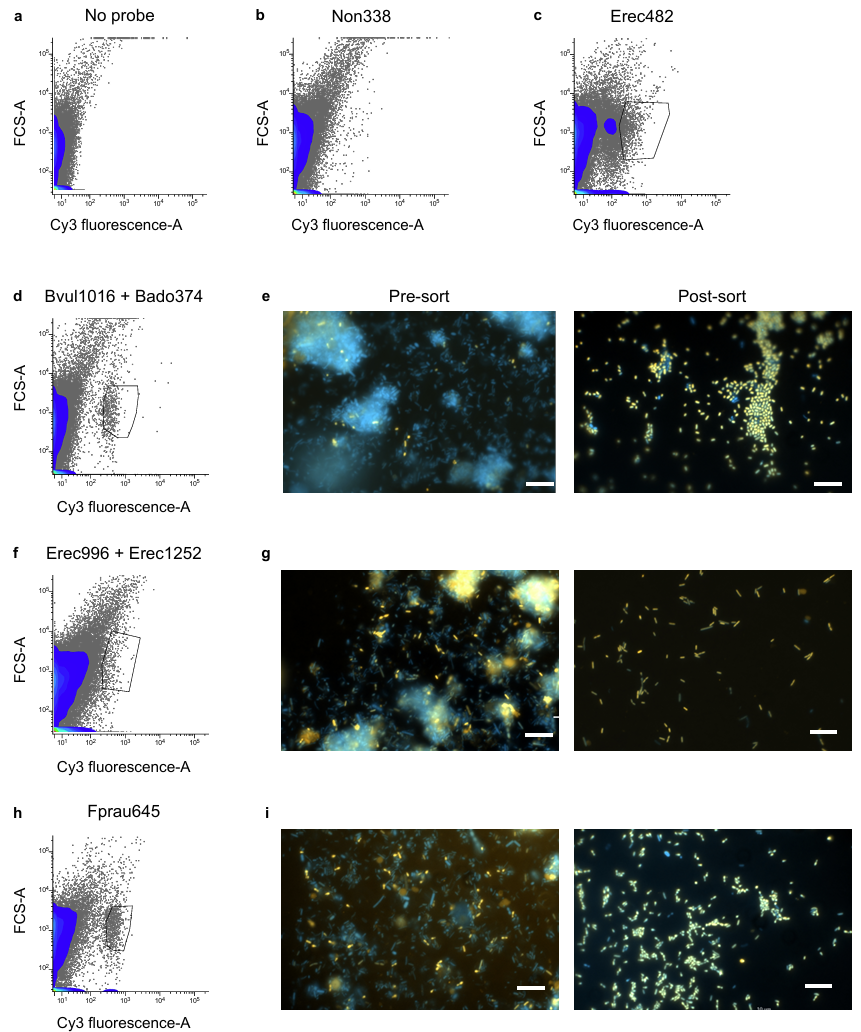

**Figure S6. FISH-FACS of DSP-fixed cells from human stool samples for metagenomic sequencing.** Representative scatter plots showing FISH-labeled cells detected by forward scatter (FSC-A) and Cy3 fluorescence-A following separate hybridization of a human fecal sample with double-labeled (DOPE-FISH) probes: **a**, no probe control, **b**, NON338, **c**, EREC482 (*Lachnospirales*), **d**, BVULG1017BADO374 (*Phocaeicola vulgatus* and *P. dorei*), **f**, EREC996/EREC1252 (*Agathobacter rectalis*), and **h**, FPRAU645 (*Faecalibacterium/Gemmiger*). The nonsense probe NON338 and a no probe control were included to assess background fluorescence. Sorting gates are indicated in plots **c-d**, **e**, and **f**. Representative composite epifluorescence images of FISH-labeled cells were recorded before (left) and after (right) Cy3-based cell sorting for three probe sets: **e**, BVULG1017/BADO374 (*Phocaeicola vulgatus* and *P. dorei*), **g**, EREC996/EREC1252 (*Agathobacter rectalis*), and **i**, FPRAU645 (*Faecalibacterium/gemminger*). Pre-sort images show microbial communities in the original human stool sample, while post-sort images demonstrate enrichment of probe-targeted cells after FISH-FACS. FISH-stained cells appear in yellow/orange. DAPI was used for DNA counterstaining. Cells only stained by DAPI appear in blue. Sorting was based on Cy3 fluorescence. Scale bars are 10 µm.
